# Maternal Wnt11b regulates cortical rotation during *Xenopus* axis formation: analysis of maternal-effect *wnt11b* mutants

**DOI:** 10.1101/2022.02.02.478872

**Authors:** Douglas W. Houston, Karen L. Elliott, Kelsey Coppenrath, Marcin Wlizla, Marko E. Horb

## Abstract

Asymmetric signalling centres in the early embryo are essential for axis formation in vertebrates. These regions, namely the dorsal morula, yolk syncytial layer, and distal hypoblast/anterior visceral endoderm (in amphibians, teleosts and mammals, respectively), require the localised stabilisation of nuclear Beta-catenin (Ctnnb1), implying that localised Wnt/Beta-catenin signalling activity is critical in their establishment. However, it is becoming increasingly apparent that the stabilisation of Beta-catenin in this context may be initiated independently of secreted Wnt growth factor activity. In *Xenopus*, dorsal Beta-catenin stabilisation is initiated by a requisite microtubule-mediated symmetry-breaking event in the fertilised egg: “cortical rotation”. Vegetally-localised *wnt11b* mRNA has been implicated upstream of Beta-catenin in this context, as has the dorsal enrichment of Wnt ligand-independent activators of Beta-catenin, but the extent that each of these processes contribute to axis formation in this paradigm remains unclear. Here we describe a maternal effect mutation in *Xenopus laevis wnt11b*.*L*, generated by CRISPR mutagenesis. We demonstrate a maternal requirement for timely and complete gastrulation morphogenesis and a zygotic requirement for proper left-right asymmetry. We also show that a subset of maternal *wnt11b* mutants have axis and dorsal gene expression defects, but that Wnt11b likely does not act through the Wnt coreceptor Lrp6 or through Dishevelled, which we additionally show (using exogenous constructs) do not exhibit patterns of activity consistent with roles in early Beta-catenin stabilisation. Instead, we find that microtubule assembly and cortical rotation are reduced in *wnt11b* mutant eggs, leading to less organised and directed vegetal microtubule arrays. In conclusion, we propose that Wnt11b signals to the cytoskeleton in the egg or early zygote to enable robust cortical rotation, and thus acts in the distribution of putative dorsal determinants rather than as a component or effector of the determinants themselves.

## Introduction

The formation of a single dorsal midline axis of bilateral symmetry in the embryo is a central event in the early development of vertebrate animals. In amphibians, this process depends on symmetry-breaking in the fertilised egg, which occurs through a process of microtubule-mediated “cortical rotation” of the egg cortex, resulting in corticocytoplasmic translocation of putative dorsal axial determinants (Gerhart, 2004; Houston, 2012; Houston, 2017; Weaver and Kimelman, 2004). A major outcome of cortical rotation is the stabilisation and nuclear localization of Ctnnb1/Beta-catenin (Beta-catenin hereafter) during the 16-to-32 cell stages (Kao and Elinson, 1988; Yang et al., 2002). This process sets up chromatin states and later gene regulatory interactions (Blythe et al., 2010) that lead to the specification of cells comprising the Spemann organiser (reviewed in Houston, 2017). The organiser ultimately elaborates axial patterning through the formation of extra-cellular gradients of BMP (and other) growth factor antagonists.

At least two non-mutually exclusive views of dorsal signalling subsequent to cortical rotation have been considered: 1) the intracellular localization of cytoplasmic activators of Wnt signalling in the absence of Wnt ligand, and 2) the dorsal enrichment of vegetally localised Wnt ligand (*wnt11b*) mRNA. The (first) intracellular localization model is supported by various studies, including: cytoplasmic ablation/transplantation experiments involving egg cytoplasm (Darras et al., 1997; Kageura, 1997; Marikawa and Elinson, 1999; Marikawa et al., 1997), the visualisation of translocating particles containing exogenous Dishevelled (Dvl) and/or Frat1/GBP fusion proteins (Miller et al., 1999; Weaver et al., 2003), and the failure of overexpressed extracellular Wnt antagonists to inhibit endogenous beta-catenin activity or axis determination (Hoppler et al., 1996; Leyns et al., 1997; Wang et al., 1997).

Also, recent work in fish and frogs has shown that deficiency in maternal *huluwa* (*hwa*; (Yan et al., 2018), which encodes a localised RNA in both species, results in ventralization. These data also included dominant-negative results suggesting that Hwa acts independently of secreted Wnt signalling, supporting the prior results in frog embryos. Hwa accumulates dorsally and likely promotes the dorsal degradation of Axin1, a key scaffold protein that facilitates beta-catenin degradation (Yan et al., 2018). More broadly, Beta-catenin is required for the establishment of the Anterior Visceral Endoderm (AVE) in mouse embryos, the requisite axis-regulating region, but evidence is accumulating that the AVE can form in the absence of secreted Wnt ligand signalling (reviewed in Houston, 2017).

The (second) extracellular signalling model was suggested by experiments showing that depletion of maternal *wnt11b* mRNA using antisense oligonucleotides (oligos) results in axial defects, with corresponding loss of early Beta-catenin target gene expression (Tao et al., 2005). These studies also indicated that Wnt11b is required extracellularly, in a complex with Exostosin glycosyltransferase 1 (Ext1) and Tdgf1 (*alias* Cripto/FRL-1) homologues, and possibly with other Wnt ligands (Cha et al., 2008; Tao et al., 2005), implying an extracellular signalling event. TALEN-mediated mutagenesis in zebrafish has ruled out a maternal role for *wnt8a* in early development (Hino et al., 2018), although the roles of other Wnts have yet to be tested in this organism.

These studies on maternal Wnt signalling need to be reconciled with more general data on Wnt/Beta-catenin signalling showing the importance of activated Wnt receptor-coreceptor complexes (Lrp6 “signalosomes”) in association with Dvl (Bilic et al., 2007). A hybrid model of maternal Beta-catenin regulation has been proposed wherein cortical rotation might enrich active phospho-Lrp6 signalosomes on the dorsal side (Dobrowolski and Robertis, 2012). However, recent data suggest that endocytosis may not be required for Wnt/Beta-catenin activation (Rim et al., 2020). And, the extent that phospho-Lrp6 signalosomes become dorsally enriched is unknown.

To re-evaluate the role of maternal Wnt11b signalling in axis formation, we used targeted genome editing to generate *Xenopus laevis* deficient in *wnt11b*, obtaining embryos lacking Wnt11b function maternally, zygotically, or both. Our data show that maternal Wnt11b signalling is required for proper initiation of dorsal gene expression, axis formation, and gastrulation morphogenesis, but that axial tissues eventually form in many cases. Additionally, consistent with prior reports, we find that loss of Wnt11b activity is associated with left-right asymmetry defects. Regulation of Lrp6 phosphorylation and Dishevelled puncta were unchanged in *wnt11b* mutant oocytes and eggs. Furthermore, and in contrast to the existing models for Wnt11b in Beta-catenin activation, live imaging experiments show that vegetal microtubule dynamics and cortical rotation are disrupted in the absence of Wnt11b function, implicating Wnt11b in the initiation or maintenance of, rather than (or in addition to) the outcome of, cortical rotation.

## Results

### Mutagenesis of *Xenopus laevis wnt11b*

CRISPR/Cas9 mutagenesis was used to create insertion-deletions (indels) in the *wnt11b*.*L* locus of the inbred *Xenopus laevis* J-strain (Session et al., 2016; TOCHINAI and KATAGIRI, 1975). These mutants were generated and raised at the National *Xenopus* Resource (NXR; (Pearl et al., 2012). We designed guide RNAs targeting the first two exons of the *wnt11b*.*L* gene on chromosome 8L (chr8L; hereafter just *wnt11b*)(**Fig. S1A, Table S1**). These guide RNAs lack homology to the paralogous *wnt11*.*L/S* genes (*alias wnt11-r*), which are located on chromosomes 2L/2S and are not expressed maternally, but share similar zygotic expression patterns to *wnt11b* (Garriock et al., 2005) (**Fig. S1B**). The homeolog (*alias* alloallele) for *wnt11b*.*S*, which would be present on chromosome 8S, is absent, a likely consequence of widespread gene loss on the S subgenome chromosomes (Session et al. 2016). *X. laevis* also lacks a duplicated *wnt11b* gene found in *X. tropicalis* (*alias xetro*.*H00536*; (Dichmann et al., 2015)). Thus, *X. laevis* is functionally diploid for the *wnt11b* gene, and Wnt11b represents the sole contribution to maternal Wnt11 protein family function.

Guide RNAs against *wnt11b* were complexed with Cas9 protein and injected into one-cell embryos obtained from matings/in vitro fertilizations of wildtype J-strain *X. laevis* to begin generating a mutant pedigree (**Fig. 1A**). A set of resulting embryos were grown to sexual maturity and females were tested for germline transmission of indels. An F0 female was identified that transmitted a 13 bp deletion (−13del; allele designation *Xla*.*wnt11b*^*emNXR*^), located near the three prime end of exon1 (sgT1 guide RNA site). This deletion is predicted to generate a frameshift mutation resulting in a premature stop codon just after the Wnt11b signal sequence cleavage site (**Fig. 1B**). Outcrossing of this ‘founder’ female to a wildtype J strain male resulted in heterozygous F1 progeny that were then intercrossed. Three sexually mature F2 females homozygous for the 13 bp deletion were initially identified, in addition to several homozygous mutant males (**Fig. 1A**). These F2 generation *Xla*.*wnt11b*^*emNXR/emNXR*^ (*wnt11b*-/-hereafter) frogs all developed normally (outwardly), indicating that zygotic *wnt11b* is not uniquely required for developmental viability.

**Fig. 1.**
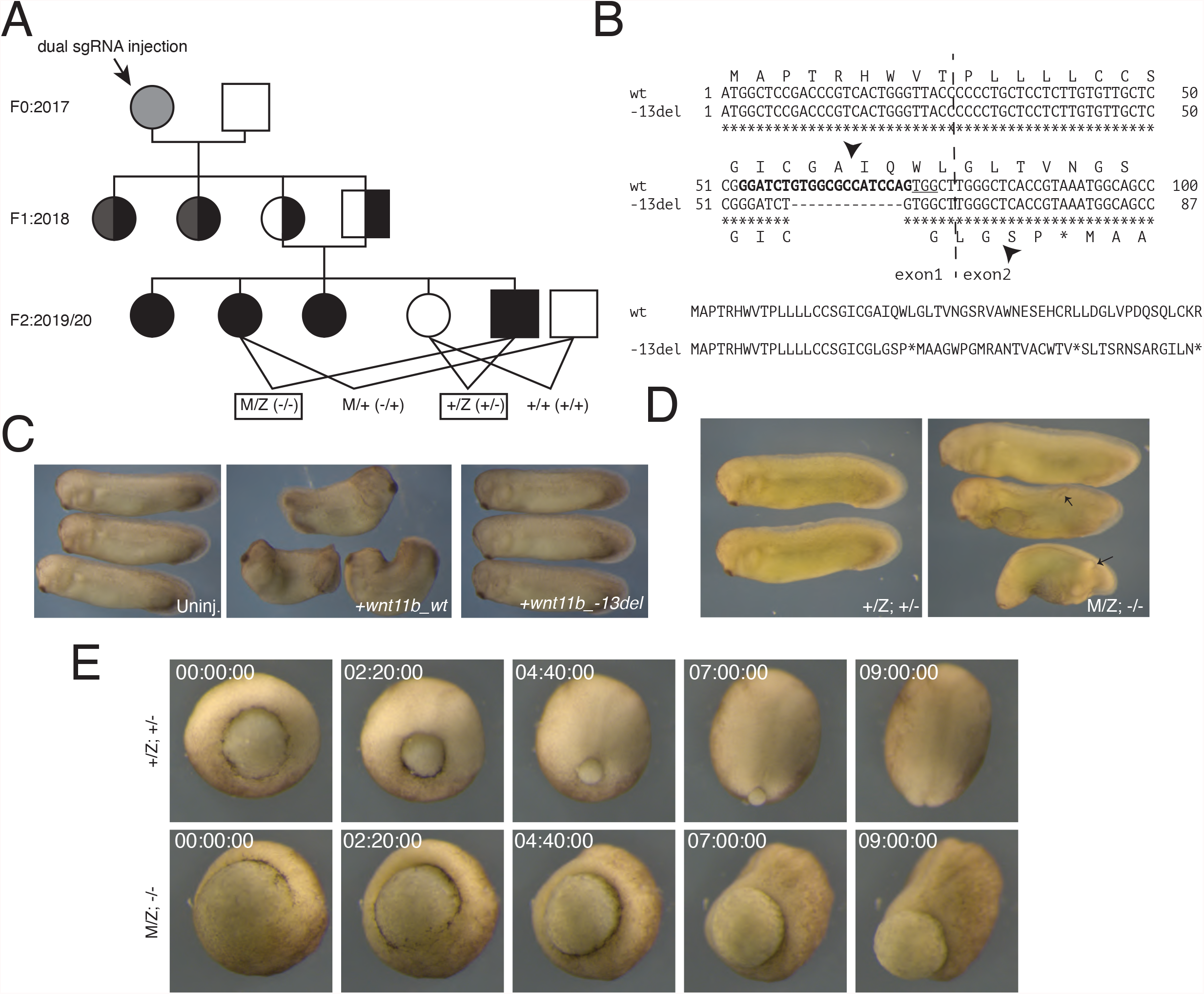
Mutagenesis of *wnt11b*.*L* in *Xenopus laevis*. (A) Genealogy of *Xla*.*wnt11b*^*emNXR*^ maternal-effect mutants. (B) Alignments of wildtype (wt) and mutant (−13del) nucleotide and predicted protein sequences. *=identical nucleotide residues (or stop codon in protein sequence). The CRISPR sgRNA target is bolded and the PAM is bolded. The dashed line represents the exon1:exon2 boundary. Arrowheads indicate the predicted signal peptide cleavage sites. (C) Phenotypes of embryos injected with *wnt11b* wildtype mRNA (*wnt11b_wt;* middle), *wnt11b* mutant mRNA (*wnt11b_-13del;* right) or left uninjected (Uninj.; left). (D) Representative images of heterozygous controls (+/Z; left) and maternal-zygotic *wnt11b* mutants (M/Z; right). Arrows indicate ectopic tailbud extensions. (E) Representative frames from time-lapse movies of a heterozygous control gastrula (+/Z; top) and a sibling maternal-zygotic *wnt11b* gastrula (M/Z; right). Time stamps indicate time from the beginning of filming (hr:min:sec).

To assess the functionality (or lack thereof) of the *-13del/emNXR* allele, we amplified and cloned the *wnt11b*.*L* coding region cDNAs from wildtype and mutant oocyte RNA by RT-PCR. Sequencing confirmed that the 13bp deletion created the predicted frameshift mutation, resulting in premature stop codons in the expressed RNA, and that it did not disrupt normal exon1-exon2 splicing (**Fig. 1B**). Additionally, we synthesised transcripts from these cDNAs in vitro and assessed their activity through overexpression in wildtype *Xenopus* embryos. In all cases (n>=30 each; two experiments), wildtype *wnt11b* induced axis shortening, consistent with known phenotypic effects of *wnt11b* injection (**Fig. 1C**; (Du et al., 1995). By contrast, overexpression of mutant *wnt11b-13del* transcripts had no effect on development and morphogenesis (**Fig. 1C**), a result further in line with this mutant allele lacking activity.

### *wnt11b* is maternally required for normal progression through gastrulation and axial morphogenesis

Homozygous *wnt11b-/-* F2 females produced fertile eggs, and homozygous mutant F2 males were similarly fertile, indicating that zygotic Wnt11b function is not required for overall germline development. To assess the role of maternal Wnt11b in development, we fertilised eggs from homozygous mutant females using sperm from either wildtype or mutant males, alongside eggs from wildtype females. Because we wished to unambiguously describe the strictly maternal effects of *wnt11b* loss-of-function, we designated the origin of the mutant allele in these crosses using a capital ‘M’ or ‘Z’ for whether the mutant allele was maternally/egg derived (‘M’) or zygotically/sperm derived (‘Z’). Thus maternal-zygotic mutants are referred to using ‘M/Z’ in this work, whereas heterozygous embryos are indicated as ‘M/+’ or ‘+/Z’, corresponding to mutant or wildtype eggs fertilised with wildtype or mutant sperm, respectively. For simplicity, most experiments focused on comparing M/Z (MZ*wnt11b-/-*) embryos with +/Z (+Z*wnt11b+/-*) embryos.

The crosses described above (**Fig. 1A**) resulted in embryos that developed normally until gastrulation. In embryos derived from mutant females (M/Z and M/+), gastrulation was consistently delayed with complete penetrance, regardless of paternal genotype (see below). Maternal mutant embryos subsequently developed a range of defects in embryogenesis (variable expressivity), ranging from severe axial truncation/ventralization, with or without open blastopores and neural tubes (spina bifida-like) and small ectopic tailbuds, to largely normal development (**Fig. 1D**; **Table S2**). We characterised the delays in gastrulation using time-lapse imaging of heterozygous (+/Z) and maternal/zygotic homozygous (M/Z) embryos. For these experiments, embryos were fertilised at the same time and imaged in the same dish (**Fig. 1E, Supplemental Video 1**). Movies were started when the control +/Z embryos were at the mid-gastrula stage (Nieuwkoop and Faber stage 10.5), the time when the majority of M/Z embryos were unequivocally delayed (**Fig. 1E**, time 00hr:00min:00sec). M/Z embryos did not initiate ventral blastopore closure until the equivalent stage 12 and most did not fully close the blastopore during the nine hours of imaging. Neural plate formation and morphogenesis commenced roughly on schedule in these abnormal embryos, suggesting a primary defect in gastrulation and not in dorsal signalling per se (**Fig. 1E**, time 07hr:00min:00sec). Similar results were seen using antisense Morpholino oligos against all *wnt11* family transcripts (Van Itallie et al., 2022), suggesting that normal expression of either maternal (of which Wnt11b is the sole contributor) or pan zygotic Wnt11 proteins is required for timely and complete gastrulation.

The formation of normal dorsal axial structures was first verified by assessing antigen tissue markers at the tailbud stage. Immunostaining against notochord and somite antigens using monoclonal antibodies (mAbs Tor70 and 12/101; (Bolce et al., 1992; Kintner and Brockes, 1984; Kushner, 1984)) showed reduced and disorganised expression in the population of mutant embryos that formed dorsal axes (**Fig. S2**). Interestingly, immunostaining against NCAM (mAb 6F11; (Lamb et al., 1993)) revealed ectopic neural differentiation both in ectopic tailbud structures and in epidermal regions throughout the embryo (**Fig. S2**). Embryos lacking visible axes lacked staining for these antigens (not shown). Thus, whereas a subset of embryos maternally deficient in *wnt11b* lack dorsal axes, many are able to form dorsal structures and even ectopic neural tissues.

### *Wnt11* family transcript expression is normal in *wnt11b* mutant oocytes and embryos

We next sought to determine the extent that the early stop codon in the *emNXR* mutant allele of *wnt11b* may have indirectly affected activities unrelated to Wnt11b protein function, resulting from either the degradation or delocalization of *wnt11b* transcripts during oogenesis. We also examined whether compensatory expression of *wnt11*.*L/S* genes (on chr2) might have occurred. We isolated oocytes and embryos from *wnt11b* homozygous mutant females and from wild type siblings and analysed the expression of RNAs localised to the vegetal cortex. RNAs representative of the three main classes of localised RNAs were examined: germ plasm, late pathway and intermediate RNAs, exemplified by *nanos1, vegt* and *wnt11b* itself (Houston, 2013). All three RNAs were expressed similarly when comparing wildtype and mutant oocytes (**Fig. 2A**), indicating that general oocyte polarisation and RNA localization occur normally in *wnt11b-/-* mutant animals.

**Fig. 2.**
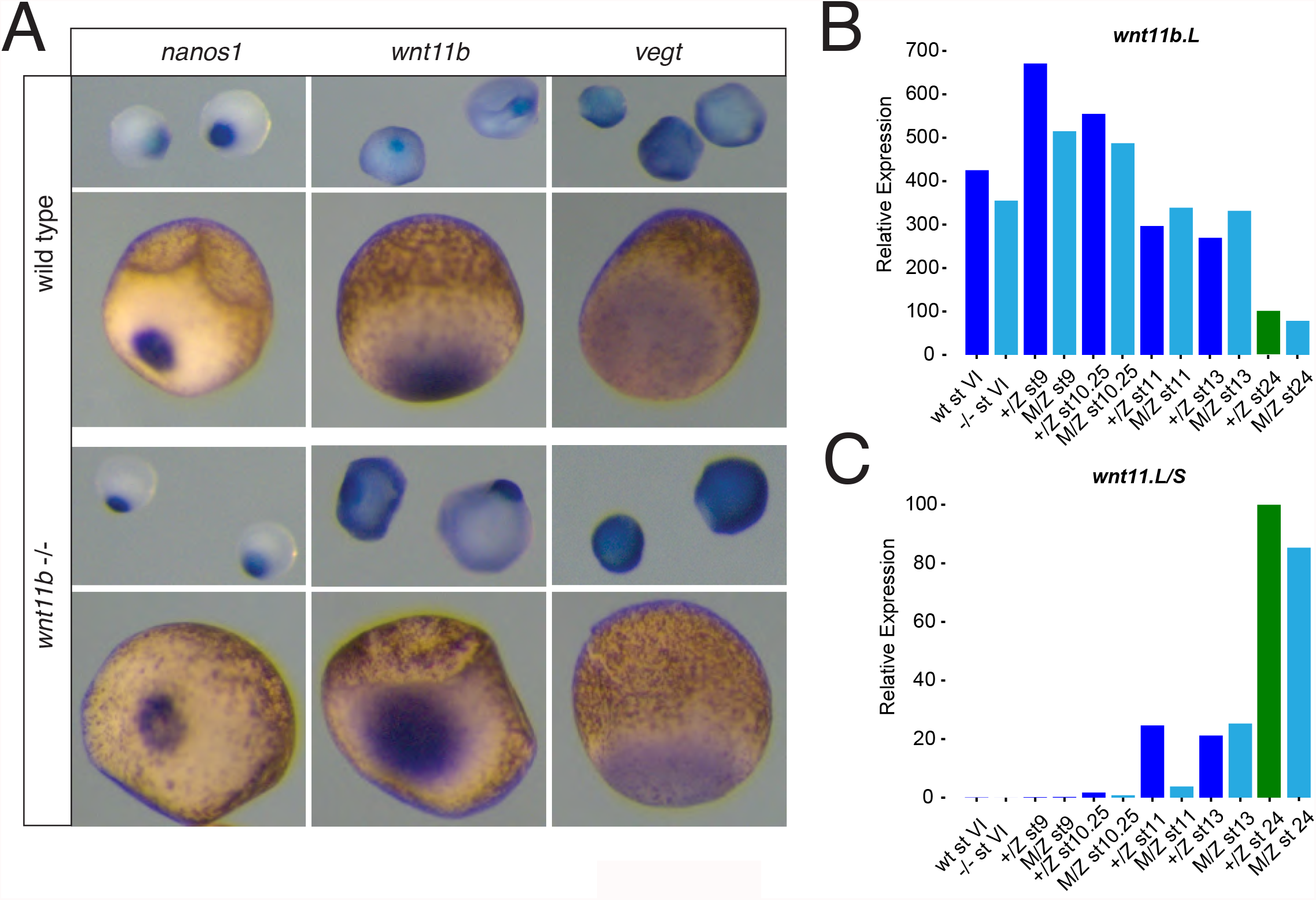
Expression of *wnt11b* in mutant oocytes and embryos. (A) In situ hybridization of *wnt11b* expression in oocytes from wildtype and homozygous mutant females (stages I-II upper row in each set; stages IV lower rows; vegetal views). (B-C) Real-time RT-PCR analysis of *wnt11b* (B) and *wnt11* (C) in wildtype and mutant oocytes (st VI) and in heterozygous and maternal-zygotic mutant embryos during development. Staging is by Nieuwkoop and Faber (1956). Bars coloured green indicate that the st24 sample was used for normalisation for relative expression; mutant samples are coloured in cyan.

We also assessed *wnt11b* and *wnt11* RNA expression in oocytes and embryos by RT-PCR. Transcripts for *wnt11b* were expressed equivalently in wildtype and *wnt11b-/-* mutant oocytes and embryos throughout development (**Fig. 2B**), suggesting a lack of nonsense-mediated decay maternally. *wnt11*.*L/S* paralogues were not ectopically expressed maternally in *wnt11b-/-* mutant oocytes or during later stages (**Fig. 2C**), although their expression was delayed around stage 11 (mid-gastrulation), likely as part of the general delay in gastrulation in these embryos (see below).

These data suggest that the variable defects seen in *wnt11b* maternal mutants are the result of loss of maternal Wnt11b protein function, and that variable up-or down-regulation of the paralogous *wnt11*.*L/S* genes or disruption of RNA localization does not generally occur and does not appear to contribute to the *wnt11b*-deficient phenotypes.

### *wnt11b* is required zygotically for left-right asymmetry

In the course of these initial studies on the role of maternal *wnt11b*, we noted that a homozygous *wnt11b-/-* F2 male used as a testis donor had reversed organ situs. Because *wnt11b* had previously been implicated in laterality signalling using Morpholino-based knockdown (i.e., loss of left-sided *pitx2c* expression; (Walentek et al., 2013)), we performed heterozygous crosses and scored resulting embryos at the swimming tadpole stage, when organ laterality is easily discernible (Yost, 1992). In a pilot experiment (47 total tadpoles), we identified three tadpoles that exhibited reversed heart orientation (heterotaxy or situs inversus; abnormal gut coiling and/or heart orientation). Genotyping revealed that all three heterotaxic tadpoles were homozygous for *wnt11b-/-*, whereas the remaining 8 homozygous mutant tadpoles and the heterozygotes and wildtype tadpoles had normal situs (situs solitus; **Table S3**), a frequency of about 25% laterality defects in homozygous F2 mutants.

Subsequent observations of surviving F3 MZ*wnt11b-/-* mutant tadpoles with normal phenotypes showed a similar frequency of heterotaxy (24-25%; n >= 50; **Table S3, Fig. 3A-B**). Examination of *pitx2c* expression at the tailbud stage showed normal expression in the left lateral plate mesoderm of heterozygous embryos (+/Z; **Fig. 3C-D**) and in about half the maternal-zygotic mutants (M/Z; **Fig. 3E-E’**). Left-sided *pitx2c* could be seen in some axis deficient embryos, suggesting left-right defects could be independent from axial induction defects (**Fig. 3E**). The remaining maternal-zygotic mutant embryos lacked or were severely reduced in left-sided *pitx2c* expression, or exhibited bilateral *pitx2c*, with weak expression on the right side (**Table S3**). These frequencies are comparable to the frequency seen in the null embryos from heterozygous matings (above), and in published studies on *wnt11b* Morpholino-injected embryos (in which zygotic, but not maternal Wnt11b, function is affected; Walentek et al. 2013). These congruences suggest that the action of Wnt11b on left-right signalling is mainly performed by zygotically expressed *wnt11b*. We did not pursue this zygotic aspect of Wnt11b’s function further.

**Fig. 3.**
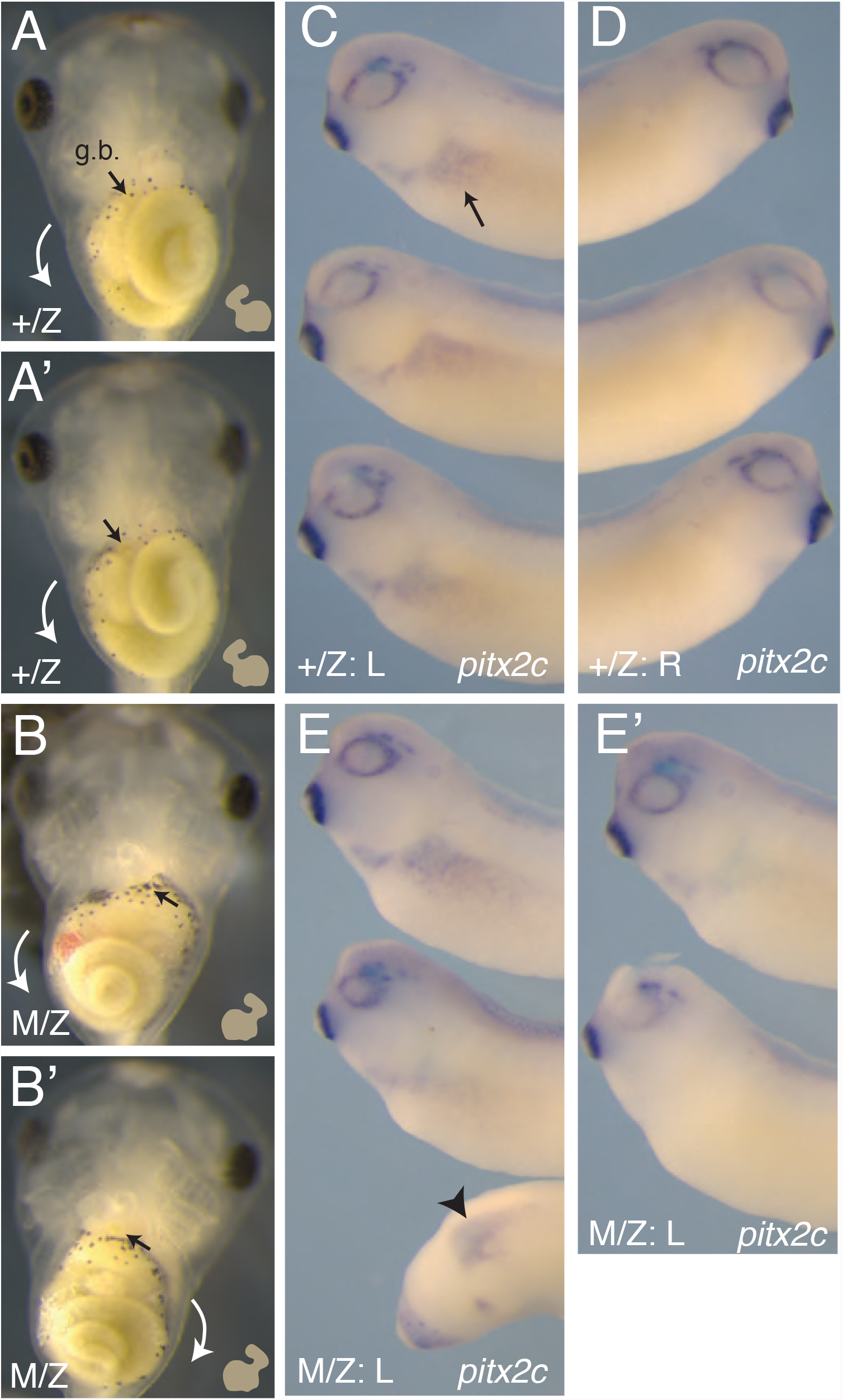
Laterality defects in *wnt11b* mutant embryos. (A-A’) Ventral views of stage 46 control heterozygotes (+/Z), showing normal lateral asymmetry. The heart shape is diagrammed (lower left), showing the ventricle positioned on the left-side and the outflow tract oriented to the right. The intestines coil counterclockwise, as shown by the arrows. (B-B’) Ventral views of maternal-zygotic mutant embryos at stage 46, showing examples of heterotaxy and *situs inversus*, respectively. The heart orientation is reversed in both examples, and the gut coils clockwise in (B’). (C) Left-sided *pitx2c* expression at stage 30 in heterozygous control embryos. (D) Absence of *pitx2c* expression on the right side of control embryos. (E-E’) Left-sided *pitx2c* expression in M/Z mutants; arrowhead indicates example of normal *pitx2c* expression in an axially truncated embryo (E) and absence of *pitx2c* expression on the left side of a subset of M/Z mutant embryos.

### Maternal Wnt11b is required for normal early dorsal-ventral gene expression

To characterise the maternal effects of *wnt11b* loss-of-function, we analysed gene expression in embryos derived from *wnt11b-/-* F2 females at the gastrula and stages, when visible phenotypes first became apparent. Consistent with visible delays in gastrulation, the expression of mesoderm and endoderm markers was reduced and delayed during gastrulation in maternally *wnt11b* deficient embryos (**Fig. 4**). Notably, mesodermal genes *myod1* (stages 12-13), and *wnt11b* itself and *wnt8a* (stage 10.5) were expressed in patterns consistent with delayed blastopore formation in maternal mutant embryos. (**Fig. 4A-L, Q**). This delay was particularly evident in delayed ventral endoderm expression of *sox17a* (**Fig. 4M, O**), whereas a general ventral fate marker, *sizzled (szl)*, was expressed in its normal spatial pattern (**Fig. 4D, F, Q**), although at elevated overall levels. The expression of dorsal and Wnt/Beta-catenin target genes were also reduced in maternal *wnt11b* mutant embryos (MZ*wnt11b-/-* and M+*wnt11b-/+*). In situ hybridization analyses of dorsal *nodal3*.*1* expression showed either reduced or absent expression in all cases, with about 50% lacking visible signal (**Fig. 4N, P**; **Fig. S3**). Similarly, in RT-PCR analyses, levels of the organiser marker *siamois homeodomain 1* (*sia1*; **Fig. 4R**) and *nodal3*.*1* (**Fig. S3**) were reduced to =< 20% of control levels (on average) in embryos derived from *wnt11b* mutant eggs (M/Z and M/+). Conversely, expression of the ventral marker *szl* was expressed at normal (low) levels at stage 9, but was elevated in maternal *wnt11b* mutant embryos at stage 10.5 (**Fig. 4S**).

**Fig. 4.**
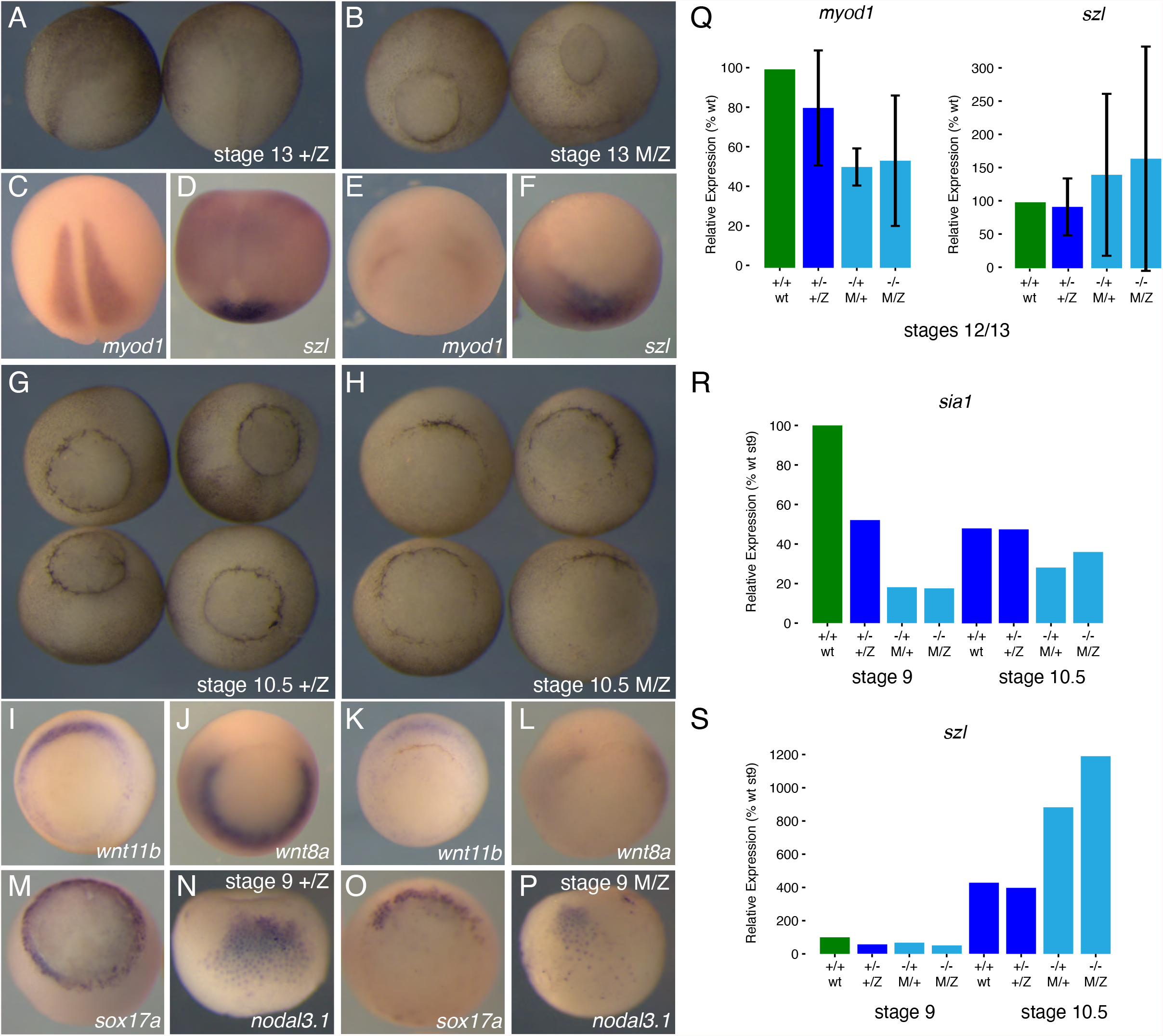
Delayed and reduced mesendodermal gene expression in the absence of maternal *wnt11b*. (A,C-D,G,I-J,M-N) Phenotypes (A, G) and in situ hybridization of mesendodermal markers (C, *myod1*; D, *szl*; I, *wnt11b*; J, *wnt8a*; M, *sox17a*; N, *nodal3*.*1*) in early neurula plate stage (A-D) and mid/early gastrula stages (G-N) in control heterozygotes (+/Z; left-hand panels). (B,E-F,H,K-L,O-P) Phenotypes (B, H) and in situ hybridization of mesendodermal markers (E, *myod1*; F, *szl*; K, *wnt11b*; L, *wnt8a*; O, *sox17a*; P, *nodal3*.*1*) in early neurula plate stage (A-D) and mid/early gastrula stages (G-N) in maternal mutant homozygotes (M/Z; right-hand panels). Dorsal, posterior views (A-F); vegetal views (G-P, dorsal towards top). (Q-S) Real-time RT-PCR analysis of *myod1* and *szl* at stage 12/13 (Q), and *sia1* (R) and *szl* at stage 9 and 10.5 (S). Bars in green indicate the samples used for normalization of relative expression; maternal mutants are coloured in cyan. Error bars in (Q) represent standard error of the mean of two biological replicates; representative experiments are shown for (R-S).

Preliminary RNA sequencing analysis showed that other mesendodermal genes (e.g., *mixer, foxc2*) and ectodermal/prospective neuroectoderm markers (e.g., *lhx5, sox2*) were either unaffected or slightly changed in mutants (up or down; **Fig. S4A-B**). Levels of *wnt5a*.*L/S* were slightly elevated in MZ*wnt11b-/-* embryos at stage 9 (**FigS4C**) and stage 10.5 (not shown). There was variability across replicate samples, consistent with the phenotypic variability (e.g., see M/Z replicate one versus replicate two in **Fig. S4B**). Additionally, analyses of differentially expressed genes showed dysregulation of apparently unrelated genes involved in metabolism, particularly at stage 9, which may be indicative of developmental delay (**Fig. S4D**). Thus, *wnt11b* is strictly, but variably, required maternally for normal dorsoventral patterning and for the normal progression through gastrulation.

To confirm the specificity of these effects, we conducted rescue experiments by injecting oocytes obtained from a *wnt11b-/-* F2 female with *wnt11b* mRNA and then fertilising these oocytes using host-transfer methods. We injected a low dose (20 pg) of transcript to avoid overstimulation of Wnt signalling. Injected and uninjected mutant oocytes were cultured for 24 hours, stimulated to mature, and then transferred to the body cavity of a wild type female. After subsequent ovulation, host-transferred eggs were fertilised with sperm from a homozygous mutant male. Embryos derived from mutant oocytes injected with wildtype *wnt11b* exhibited somewhat restored levels of organiser genes and underwent normal gastrulation (**Fig. S5; Table S2**).

### Phospho-Lrp6 and Dvl regulation are normal in *wnt11b* mutant eggs

Lrp6 activation has been hypothesised to initiate Beta-catenin stabilisation during axis formation, either in oocytes, or as part of signalling endosomes translocated during cortical rotation. We therefore sought to determine the patterns of Lrp6 phosphorylation during early development and to identify the extent to which these patterns were dependent on maternal Wnt11b. We assessed the state of Lrp6 activation in oocytes, eggs and early embryos by immunoblotting against a phospho-epitope of Lrp6, S1490, that is elevated by Wnt ligand binding and essential for the response to Wnt signals (Tamai et al., 2004; Zeng et al., 2005). To generate a more robust and reproducible signal, we injected oocytes with ‘trace’ amounts of exogenous mouse *Lrp6* mRNA (∼20-50 pg).

In wildtype samples, S1490 Lrp6 phosphorylation was absent in stage VI oocytes but stimulated in progesterone-treated/mature oocytes (**Fig. 5A**). Endogenous phospho-Lrp6 was weakly detected in mature oocytes but not in untreated stage VI oocytes. Furthermore, this phosphorylation was transient and was downregulated following either prick activation or normal fertilisation of host-transferred eggs. Notably, phospho-Lrp6 was substantially reduced by 60 minutes post-fertilization, when cortical rotation would be occurring, and remained absent in cleavage stage embryos (**Fig. 5B**), when Beta-catenin stabilisation would be occurring. Phenotypic analysis of sibling embryos confirmed that the low doses of *Lrp6* did not induce axial duplication/dorsalization when expressed in oocytes, whereas higher doses would do so (**Fig. S6**), indicating that the trace amounts of mRNA used are below the threshold for strongly activating Beta-catenin under these conditions.

**Fig. 5.**
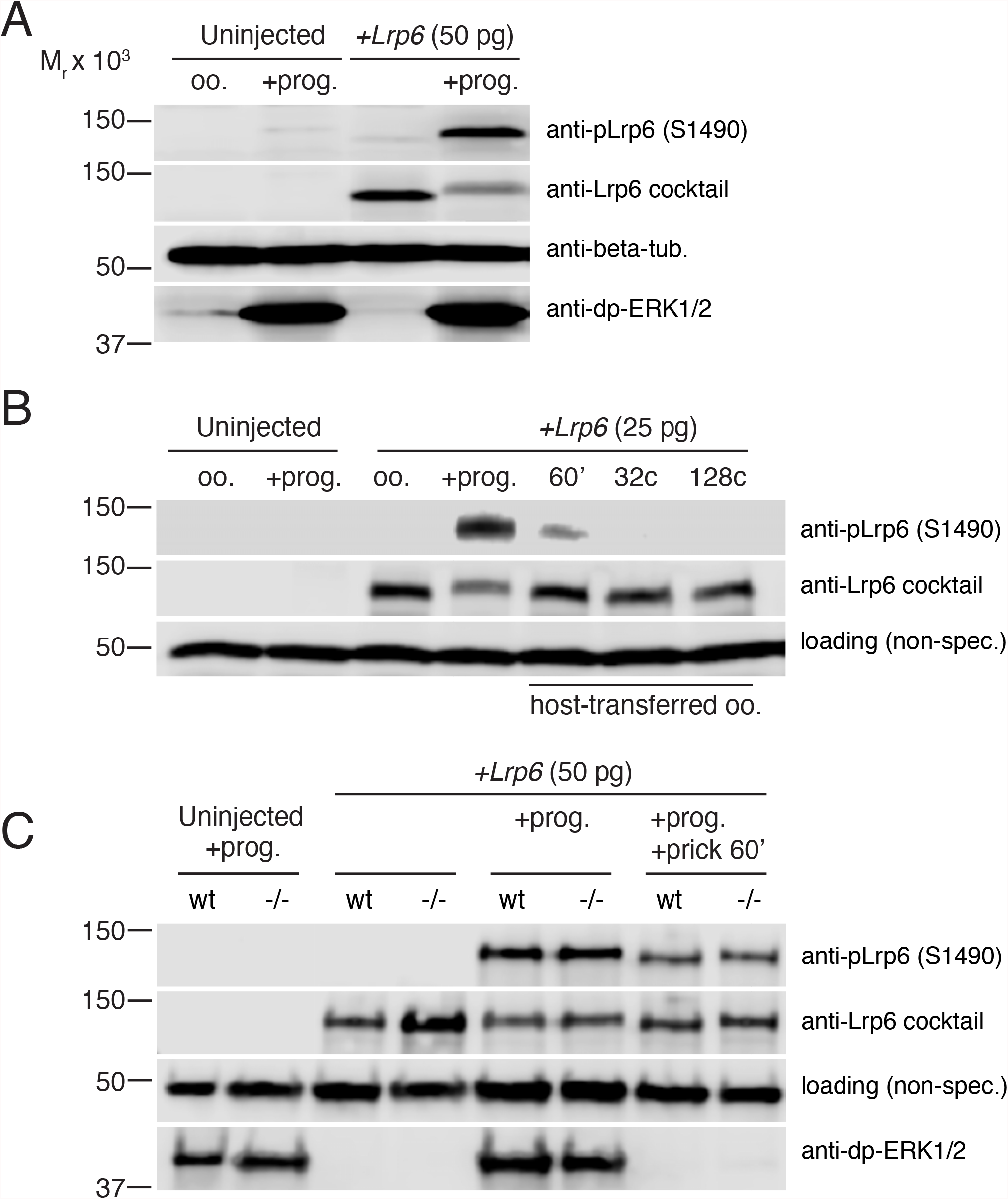
Wnt11b does not regulate the activation of Lrp6 phosphorylation during oocyte maturation. (A) Immunoblotting of control oocytes and oocytes injected with mouse *Lrp6* mRNA, untreated (oo.) or treated with progesterone (+prog.). The same blot was probed using anti-pLrp6 (S1490) and two anti-Lrp6 mAbs (anti-Lrp6 cocktail). Tubulin and di-phospho-ERK were used as loading controls. (B) Immunoblotting of oocytes as above; a subset of *Lrp6*-injected oocytes were fertilised by host-transfer and blotted against anti-pLrp6 (S1490) and two anti-Lrp6 mAbs (anti-Lrp6 cocktail). A non-specific (non-spec.) band was used to assess equal loading. (C) Immunoblotting of wildtype (wt) and homozygous mutant female (−/-) oocytes, treated with progesterone, without or with prick activation (+prick 60’). Relative migration (M_r_) against protein molecular weight standards are shown on the left. Bands were visualised by chemiluminescence.

Analysis of Lrp6 phosphorylation in *wnt11b* mutant oocytes and eggs showed that the patterns of phospho-S1490 stimulation in eggs and subsequent downregulation were identical to wildtype samples. These data suggest that Lrp6 is phosphorylated during oocyte maturation in a Wnt11b-independent manner, and that phospho-Lrp6 disappears before cortical rotation is complete and before the critical period for Beta-catenin stabilisation (i.e., the 16-to-32-cell stages).

We also tested the working hypothesis that Wnt11b might regulate the formation or activity of Dvl puncta, which are thought to act as dorsal determinants in *Xenopus* (Dobrowolski and Robertis, 2012; Miller et al., 1999), potentially in association with activated Wnt co-receptor Lrp6 (“signalosomes”; (Dobrowolski and Robertis, 2012; Miller et al., 1999)). We injected transcripts encoding Dvl2-GFP into both wildtype and *wnt11b-/-* mutant oocytes. In both cases we observed numerous Dvl2-GFP puncta by live epifluorescence microscopy (**Fig. 6**). We also had hoped to examine the behaviour of these puncta during cortical rotation. However, we instead found that Dvl2-GFP puncta disappeared following oocyte maturation and did not reappear after prick-activation in both wildtype and *wnt11b-/-* mutant oocytes (**Fig. 6A-F**). Immunoblotting against GFP showed comparable levels of fusion protein expression, indicating that the loss of puncta is not the result of Dvl protein degradation (**Fig. 6G**). Additionally, we tested the propensity of Dvl2 puncta associate with endosomal structures in *Xenopus* oocytes but we failed to observe co-localization of Dvl2-mCherry with exogenous endosome markers Rab5- and Rab11-GFP in coinjected wildtype oocytes (**Fig. 6H-I**).

**Fig. 6.**
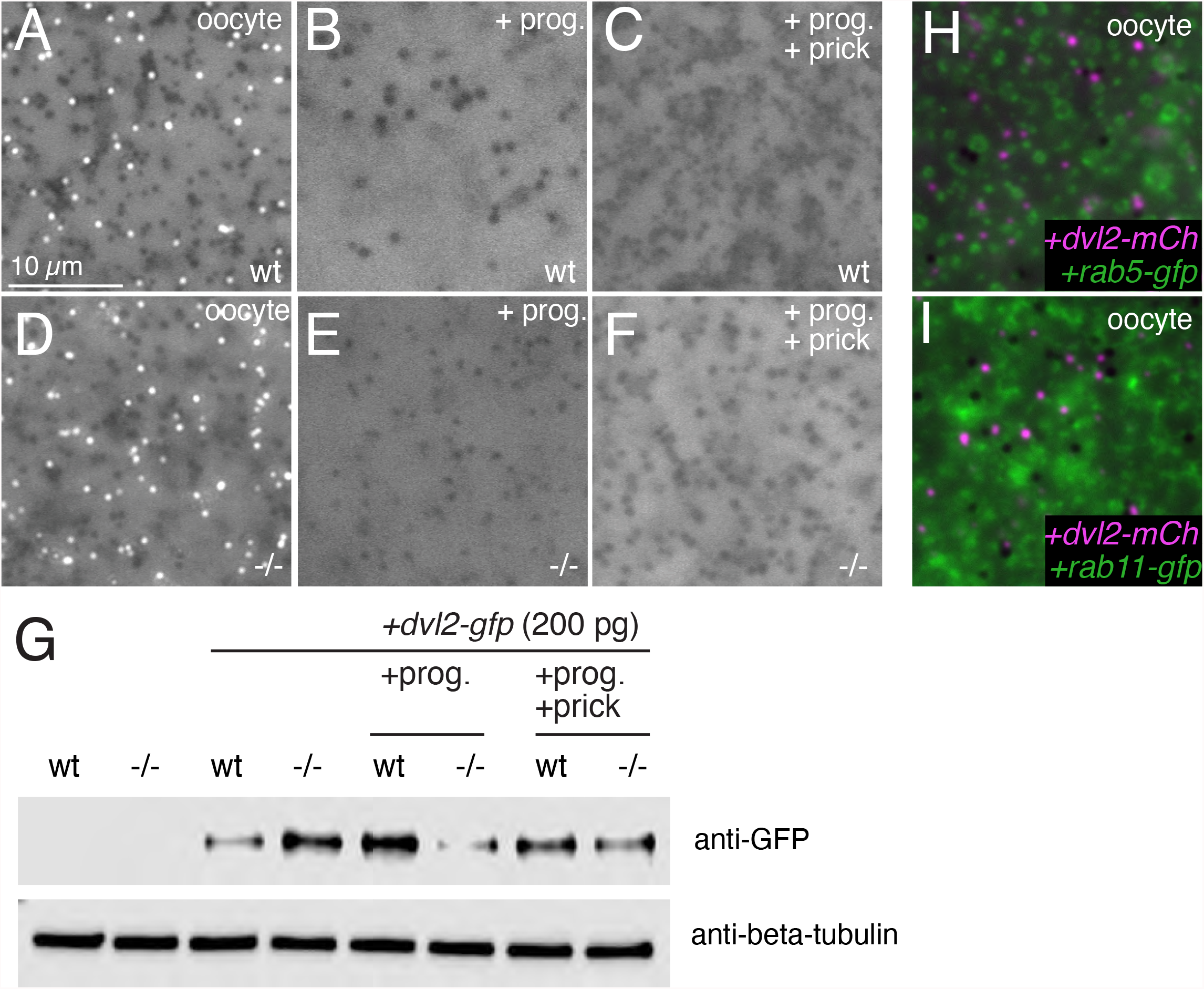
Wnt11b is not required for exogenous Dishevelled puncta regulation. (A-C) Control wildtype (wt) oocytes injected with *dvl2-gfp* and incubated untreated (A) or treated with progesterone (2 µM overnight, B) or treated and prick-activated for 60 minutes (+prick, C). (D-F) Homozygous mutant oocytes treated with or without progesterone and prick-activation. Scale bar = 10 µm. (H, I) Control oocytes injected with either *rab5-gfp* (H) or *rab11-gfp* (I) with *dvl2-mcherry*. (G) Immunoblotting of Dvl2-GFP (anti-GFP) in wildtype (wt) and mutant (−/-) oocytes treated with or without progesterone and prick-activation. Beta-tubulin was used as a loading control.

These data show that Lrp6 is phosphorylated (activated) during oocyte maturation independently of Wnt11b function. This phosphorylation is downregulated after egg activation/fertilisation, and although reduced and present during cortical rotation, becomes undetectable by the cleavage stages, when Beta-catenin stabilisation is required to occur. In contrast, Beta-catenin levels tend to mirror phospho-Lrp6 levels in cultured cell experiments using exogenous Wnt stimulation (Kim et al., 2013; Li et al., 2012). Also, these data show that large aggregates/puncta of Dvl are found in a complementary pattern to Lrp6 phosphorylation and do not localise with endosomal markers in oocytes. Thus, these results generally fail to support models invoking dorsally localised signalosomes in axis specification, although it is possible that a subset of dorsally localised complexes with smaller Dvl oligomers (Ma et al., 2020) are present, but below the limit of detection of methods used here.

### Maternal Wnt11b regulates cortical microtubule assembly and cortical rotation

Because *wnt11b* mutant embryos showed considerable variability with regard to dorsal gene expression and axis formation despite the presence of Lrp6 phosphorylation, we considered whether Wnt11b might be required for the proper distribution of so-called dorsal determinants rather than their generation. To test this idea, we first assessed general microtubule organisation in eggs derived from *wnt11b-/-* homozygous mutant females. Eggs from wildtype and mutant females were fertilised (with sperm from wildtype or mutant males) and fixed in methanol during cortical rotation, around 60 and 80 minutes after fertilisation. Immunostaining against Beta-tubulin revealed that zygotes lacking maternal *wnt11b* (regardless of sperm genotype) exhibited disorganised microtubules (**Fig. 7B-B’, Table S4**) that remained in a loose/sparse network and did not form parallel arrays. Control eggs invariably developed well-formed parallel microtubule arrays at both time points (**Fig. 7A-A’, Table S4**). Similar results were seen in prick-activated eggs derived from in vitro-matured oocytes from wildtype and mutant females (**Table S4**).

**Fig. 7.**
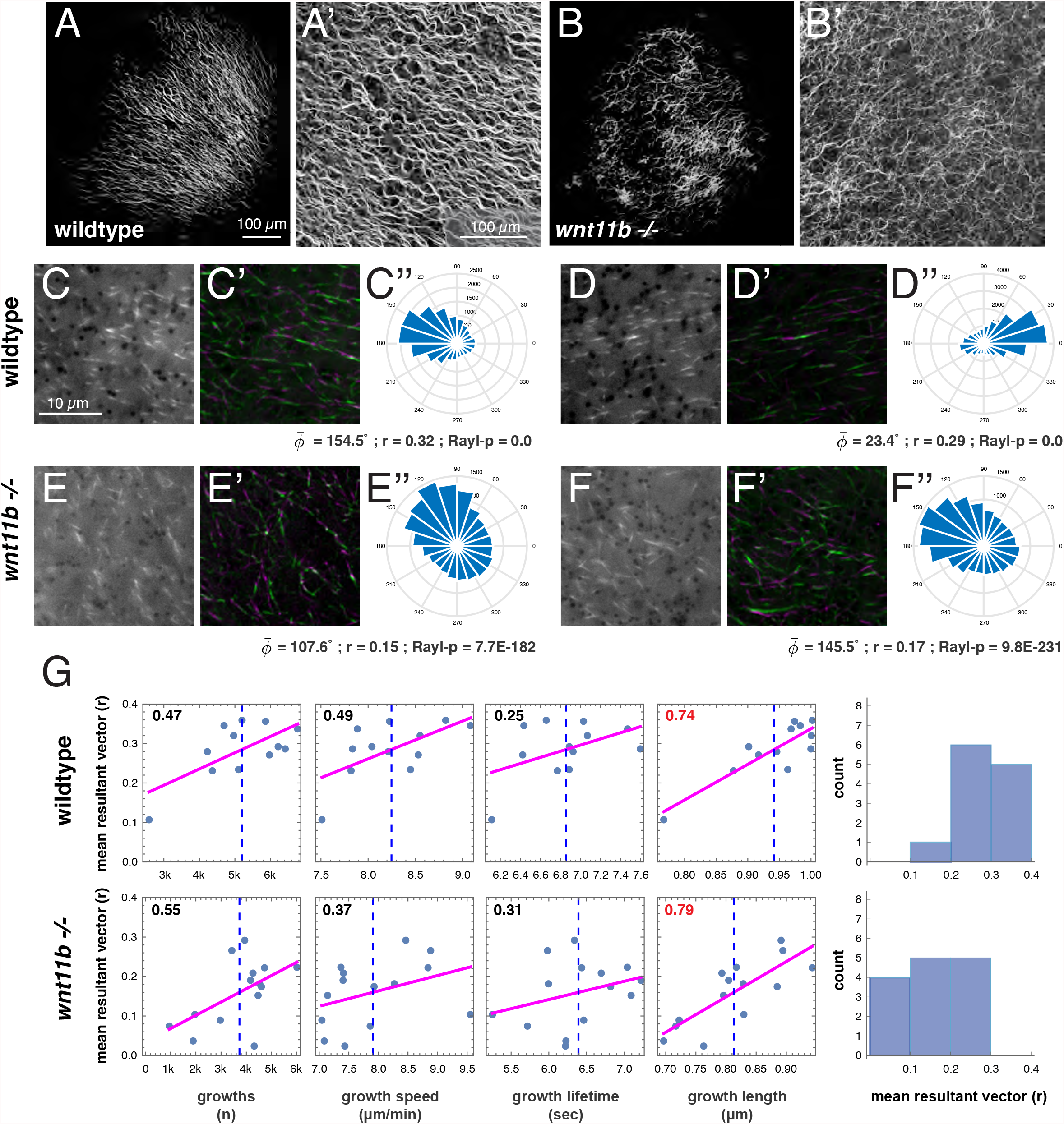
*wnt11b* is required for microtubule alignment during cortical rotation. (A-B’) Immunostaining of Beta-tubulin in fertilised wildtype (A-A’) or *wnt11b* (B-B’) mutant zygotes, fixed at 60 minutes post-fertilization. Shown are tiled images of the vegetal surface (A, B) and higher magnification confocal images (A’, B’). (C-F) Representative single frames from time-lapse movies of prick-activated wildtype (C-C’, D-D’) and *wnt11b* mutant oocytes (E-E’, F-F’) injected with *eb3-gfp* mRNA. (C’-F’) Microtubule motion is depicted by averaging frames 1-5 and frames 7-11 and merging pseudo-colored images green and magenta, respectively. (C’’-F’’) Angle histogram plots showing individual plus end track directionality (degree) per bin per two minute movie. Circular statistics for each movie are listed; 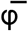(phi bar) = mean angle, r = mean resultant vector length, Rayl-p = p-value of Rayleigh test for circular uniformity. (G) Correlation plots of r versus dynamic parameters and histograms of r. Magenta lines show the least squares fit line, the blue dotted line indicates the sample mean, correlation values are upper left (red colour indicates statistical significance). The rightmost panels show histograms of r values obtained for each sample. An r > 3.0 is empirically significant in this context. Please note differing scales in correlation plots. The scale bars in (A, A’, 100 µm) apply to (B, B’), respectively. The scale bar in (C, 10 µm) applies to (C’, D, D’, E, E’, F, F’).

To more accurately measure and quantify the effect of maternal Wnt11b deficiency on microtubule dynamics and on cortical rotation, we performed live imaging of microtubule plus end dynamics. We isolated wildtype and *wnt11b-/-* mutant oocytes and injected *eb3-gfp* mRNA as a microtubule plus-end marker. These oocytes were treated with progesterone in vitro to induce maturation and then prick activated. Live imaging and analysis was performed as described (Olson et al. 2015) using oocytes, eggs, and prick-activated eggs undergoing cortical rotation (at 60 minutes post-fertilization).

Plus end dynamics in both oocytes and progesterone-treated oocytes (eggs; n=8 each) were comparable to previous reports (Olson et al. 2015) in both wildtype and mutants. Although *wnt11b-/-* mutant samples tended to have more plus end growths in oocytes with fewer plus end growths in eggs, and more rapid growth speed (**Table S5**) compared to wildtype cases, none of the differences between groups at these stages were sufficient to reject statistical non-significance. Movies of activated eggs in wildtype control samples during cortical rotation (n=12) revealed numerous clearly aligned plus-end growths, indicative of robust cortical rotation (**Fig. 7C-D**). Plus-end tracking analyses identified significant concentration around a mean angle 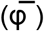 in each case, using the Rayleigh test for circular uniformity (Batschelet, 1981; Berens, 2009). Additionally, we previously established a mean resultant vector length of r = 0.3 as an empirical measure of directionality associated with the establishment of cortical rotation (Olson et al., 2015). This value was chosen because the 95% confidence intervals around the mean angle become asymptotically minimal as r approaches this magnitude. Wildtype activated eggs exhibited an average mean resultant vector length of 0.3, with over half the r values above this number (**Fig. 7G**). Only one sample registered below 0.1, a mean vector length indicative of poor directional organisation (where r=0 would be completely undirected, and r=1 would be uniformly directed on one angle).

Movies of *wnt11b* mutant eggs during cortical rotation (n=14) revealed considerable variability in plus-end growth (**Fig. 7E-F**), in line with the variability seen in anti-tubulin immunostaining and in axial development. Some eggs exhibited apparently normal directional growth, whereas others showed randomly directed, disorganised growth. Curiously, plus-end tracking analyses showed none of the mutant eggs exhibited a mean resultant vector length > 0.3, despite clearly directional growth direction in some cases. Several cases reached a magnitude of r∼0.25, but many were much lower, resulting in a lower average mean vector length (0.16 compared to 0.29 in controls; **Fig. 7G, Table S5**).

A comparison of microtubule growth parameters during cortical rotation showed that all dynamic parameters were more variable (i.e., higher variances) in the mutant eggs (**Fig. 7G**). Mean growth speed was similar in both wildtype and mutant eggs at 60 minutes post-activation, and activated *wnt11b* eggs exhibited fewer plus-end growth events in the cortex, as well as shorter growth lifetime and lengths, although only the latter difference was significant enough reject a null statistical hypothesis (**Table S6**). Plus end growth speed was also generally correlated with more robust directionality. However in one mutant sample with higher than average growth speed, directionality was compromised (9.5 µm/min, and r=0.1), suggesting a possible limit to this relationship (**Fig. 7G**).

Additional analyses of growth parameters showed significant correlation of the mean resultant vector length with plus end growth length in both wildtype samples and mutant samples (ρ=0.74 and 0.79, respectively; **Fig. 7G**). Other correlations between mean vector and growth numbers, speed and lifetime were also similar between mutant and controls, but weaker (ρ =< 0.6, p > 0.01), as were correlations of the parameters with each other (data not shown). However, correlation is a relatively weak measure of association, detecting only linear dependency independent of magnitude, and non-linear or other complex relationships might exist.

Together, these data show that the earliest detectable consequence of maternal *wnt11b* deficiency is an overall reduction in the overall organisation and alignment of vegetal cortical microtubules during cortical rotation, leading to less ‘robust’ directional orientation. This disruption is coincident with shorter growth length of microtubule plus-ends in the fertilised/activated egg. The data do not address the extent that this effect is owing to direct regulation of microtubule dynamics at the time of cortical rotation or to a more general effect on the cytoskeleton or other cellular structures or organelles following oocyte maturation.

## Discussion

We have used CRISPR/Cas9 technology to generate a loss-of-function mutation in *Xenopus laevis wnt11b*.*L*, the only maternally expressed member of the *wnt11* family in this species. Our analyses of these mutant embryos revealed a novel role for maternal Wnt11b as a permissive factor (at least) enabling robust microtubule-mediated cortical rotation and axis induction. In addition, we find that maternal Wnt11b is necessary for normal gastrulation morphogenesis, whereas zygotic Wnt11b is necessary for normal left-right asymmetry. Both of these latter findings are in line with the more traditional views of Wnt11 function obtained through antisense-mediated knockdown and dominant-negative experiments. The extent that Wnt11b might act through a similar molecular activity in all these different contexts is unknown. Wnt11b had been previously suggested to represent the main dorsalizing determinant upstream of Beta-catenin activation in *Xenopus*, whereas our results suggest that its role in this process is likely indirect.

Using targeted CRISPR/Cas9 mutagenesis of *wnt11b*.*L*, we generated a 13 base pair deletion near the end of exon1, which resulted in an early in-frame stop codon that was inherited through the germline. This deletion disrupts the normal signal peptide cleavage site and creates a premature stop codon two amino acids after a new (suboptimal) cleavage site. Thus, the mutant *wnt11b*.*L* locus would generate at most a two amino acid peptide (Serine-Proline). Several other in-frame stop codons are also generated downstream of the initial stop codon, and none of the potential peptides generated match known proteins. Based on these sequence predictions and the observation that mRNA encoding the mutant transcript lacks activity in over-expression assays, we interpret our results under the premise that this 13 base pair deletion nonsense mutation represents a null allele. Further genetic analysis, such as failure to complement deletion alleles would more conclusively demonstrate nullness, but this would require additional mutant lines.

In some contexts, premature stop codons can cause transcript degradation through nonsense-mediated mRNA decay. This mechanism does not seem to occur in the case of the -13bp deletion, which is consistent with the position of the premature stop codons at the 5’end of the transcript (Dyle et al., 2020). We examined the persistence of *wnt11b* RNA using both RT-PCR and in situ hybridization, and both assays showed normal (or slightly lowered) mRNA levels. Studies in zebrafish and mouse embryos have identified a genetic compensation response of related genes that is triggered by nonsense-mediated mRNA decay (El-Brolosy et al., 2019; Ma et al., 2019). In the absence of nonsense-mediated decay, we would also predict an absence of genetic compensation. Although our data do not directly address this phenomenon, we do note that the related *wnt11*.*L/S* genes show reduced, rather than elevated expression, a result inconsistent with compensation.

Nonetheless, also not addressed in this study, a structural or regulatory role may exist for *wnt11b* transcript, based on oligo-mediated transcript degradation experiments in host-transferred embryos (Tao et al., 2005), which exhibit a more severe ventralization phenotype. Other localised RNAs have non-coding roles in cytoskeletal organisation in Xenopus (Kloc, 2009), including *plin2* (*alias fatvg*), which is required for microtubule organisation and cortical rotation, hence axis formation (Chan et al., 2007).

It has long been problematic to reconcile and incorporate various embryological, cell biological and biochemical studies on Wnt signalling and early development into a coherent understanding of the initiating steps in axis specification in *Xenopus* (or indeed in any vertebrate embryo). The data presented here allow several refinements to current models. First, our data suggest that *wnt11b* mRNA is unlikely to be the major axis-inducing agent enriched dorsally by cortical rotation because axis signalling can occur in its absence. High-resolution transcriptomic analyses have also generally failed to identify significant dorsal enrichment or polyadenylation of *wnt11b* (Domenico et al., 2015; Flachsova et al., 2013). Second, we find that Wnt11b signalling is not required for Lrp6 phosphorylation, which is activated during oocyte maturation and then attenuated during egg activation/fertilisation. The presence of phospho-Lrp6 in the egg and its diminution following activation had been noted by Davidson et al. (2009); our observations support this idea, but also show that phosphorylation is stimulated during oocyte maturation. Third, Dvl protein forms visible puncta in oocytes but not in eggs, a pattern complementary to phospho-Lrp6 (and unchanged in *wnt11b* mutants), suggesting that these puncta (as traditionally viewed) are likely unrelated to Lrp6 activation or signalosomes. Furthermore, the disappearance of visible Dvl puncta during oocyte maturation indicates that these (large) aggregates are unlikely to be Beta-catenin-stabilising agents transported by cortical rotation either.

Our data do not rule out a small domain of persistent phospho-Lrp6 and/or small Dvl oligomers at the cell surface (à la Ma et al., 2020), which might be below the limits of detection in our assays. It is not clear to what extent small Dvl oligomers might behave differently than the larger ‘puncta’ in this context however. Other *wnts* are not highly expressed maternally, and with our present work, in conjunction with recent evidence for a cytoplasmic role for Huluwa in fish and frogs (Yan et al., 2018), the preponderance of evidence is accumulating that direct Beta-catenin stabilisation in the early morula is Wnt ligand-independent. Intriguingly, a growing body of evidence shows that regulation of Beta-catenin in the mammalian AVE occurs independently of secreted Wnt ligand signalling, but does involve Tdgf1, as is the case in frogs as well (reviewed in Houston, 2017).

One implication of these findings is that Wnt11b would be signalling in para/autocrine fashion, acting on the single-cell egg to modulate microtubule dynamics (directly or indirectly). Recent proteomic data support the idea that Wnt11b protein expression is elevated during oocyte maturation (Van Itallie et al., 2022; Peshkin et al., 2019 - visualised on Xenbase) and other studies indicate active translation in the egg (Michael D. Sheets, personal communication). This pattern is mirrored by Fzd7, a Wnt11 family receptor, suggesting that this signalling pathway becomes active in the egg.

A large body of work has demonstrated the importance of Wnt/PCP signalling mediated by Wnt11 and other ligands in the regulation of gastrulation and axial morphogenesis through the control of convergent extension and other cellular behaviours (Solnica-Krezel, 2005). In this regard, Wnt11-Frizzled7 (Fzd7) signalling is thought to regulate cortical actin/actomyosin cytoskeletal dynamics (Huebner and Wallingford, 2018), resulting in either increased or decreased cell adhesion/cohesion depending on context (Dzamba et al., 2009; Heisenberg et al., 2000; Kraft et al., 2012; Ulrich et al., 2003; Ulrich et al., 2005; Winklbauer et al., 2001; Witzel et al., 2006). Our microtubule data show that Wnt11b signalling can have an effect on the cytoskeleton independently of controlling cell motility or cell adhesion (the egg is a single cell). *Wnt11b-/-* mutant eggs might thus serve as a useful system in which to study the Wnt/PCP or Wnt/Calcium regulation of cytoskeletal dynamics without confounding feedback effects resulting from changes in cell adhesion and or polarity.

In addition to signalling in the egg, our data also show that maternal Wnt11b is essential for blastopore closure and other cell behaviours during gastrulation, in this context potentially through regulation of actomyosin and convergent extension (see above). Maternal *wnt11b* compensates for zygotic loss of *wnt11b* (in F2 homozygous mutants derived from wildtype females), and maternal mRNA injection of *wnt11b* is sufficient to rescue the M/Z phenotype. MZ*wnt11b-/-* mutants exhibit a delay in gastrulation, suggesting that other ligands (e.g., *wnt5* homologues) or signals may eventually partially compensate. We note that gastrulation in MZ*wnt11b-/-* embryos does seem to progress once zygotic expression of *wnt11 (wnt11r*) reaches normal levels (∼ stage 13) even though *wnt5* homologues are expressed before this time. However, double Morpholino-based knockdown of Wnt11b/Wnt11 results in similarly delayed gastrulation, with abnormal extension of the archenteron (Van Itallie, et al., 2022). Thus, low levels of Wnt11 family function (whether through delayed *wnt11* expression in M/Z mutants or through residual protein in knockdowns), with or without Wnt5 function, might be sufficient for ultimate closure of the blastopore. Alternatively, Wnt5 proteins could be sufficient for gastrulation to progress in the absence of Wnt11 family function, but with different (slower) kinetics. Future work, enabled by the *wnt11b-/-* mutants described here, will help distinguish these possibilities.

Maternal Wnt/PCP components (*dvl2/3, fzd7, ptk7, gpc4* and *vangl2*) are similarly required for full convergent extension in zebrafish embryos and suggest a conserved role for Wnt signals near the onset of gastrulation (Jussila and Ciruna, 2017; Xing et al., 2018). Maternal Wnt11b could regulate general cell adhesion or cell behaviours prior to gastrulation, such as those occurring in the vegetal endoderm cells (e.g., vegetal rotation, separation behaviour; (Wen and Winklbauer, 2017; Winklbauer and Schürfeld, 1999; Winklbauer et al., 2001), and thus could potentially affect convergent extension or other morphogenetic movements indirectly. Loss of Wnt11b could also alter convergent extension indirectly by altering (elevating) BMP signalling and/or expression, which is inversely correlated with convergent extension (Myers et al., 2002) or through regulation of Nodal3.1-FGFR1 signalling (Yokota et al., 2003). A more extensive analysis of signalling interactions and perigastrular cell behaviours in *wnt11b* mutants would help distinguish these possibilities.

The genetic studies presented here also support the idea of a strictly zygotic role for Wnt11b in the establishment of left-right asymmetry in *Xenopus* (Walentek et al., 2013). F2 homozygous mutants for *wnt11b* derived from heterozygous crosses undergo normal axis formation and gastrulation and yet a subset undergo altered left-right asymmetry (∼25%). A similar proportion is seen in MZ*wnt11b-/-* homozygotes. Taken together with the Morpholino-based observations of Walentek et al. (2013), which would affect only zygotic Wnt11b, these data suggest that left-right signalling is a function of zygotic Wnt11b. However, this must be only a biassing signal because roughly half the maternal/zygotic mutants develop reduced or bilateral *pitx2c* expression in the left lateral plate mesoderm – a condition which itself randomises asymmetry, thus potentially accounting for the 25% incidence of abnormal asymmetry.

In the context of left-right asymmetry, prior evidence suggests that Wnt11b controls cilia polarisation and cell shape in the ciliated cells of the posterior notochord/gastrocoel roof plate to establish leftward fluid flow (Schweickert et al., 2007; Walentek et al., 2013). Inhibition of “nodal flow” results in randomised left-right asymmetry in frogs and mice (Nonaka et al., 1998; Schweickert et al., 2010); we would therefore expect abnormal cilia and reduced fluid flow in *wnt11b* mutants. Wnt/Calcium signalling was implicated in this aspect of Wnt11b signalling, based on comparative inhibitor studies (Walentek et al., 2013) and this branch of the Wnt network was implicated in tissue separation behaviour as well (Winklbauer et al., 2001). It will thus be interesting to determine the extent that Wnt11b-mediated Wnt/Calcium signalling underlies the multiple roles of Wnt11b during development.

Other proposed functions of Wnt11b during development were not indicated by the genetic data presented here, including roles in pronephros, heart and neural crest development. It is possible that *wnt11* or other functionally related *wnts* can compensate in these contexts, or that these defects might be secondary to alterations in gastrulation caused by the various loss-of-function reagents.

Overall, it is hoped the development of maternal-effect *wnt11b* mutants (the first such engineered maternal mutation in *Xenopus*) will be valuable for studying Wnt signalling activities within individual cells, as well as in conserved gastrulation movements and cell polarity signalling at the genetic level. Wnt11b mutants could be combined with other tools commonly used in Xenopus embryology including Morpholio/oligo-or F0 CRISPR-mediated gene knockdowns to further our understanding of diverse processes in early vertebrate development.

## Materials and methods

### *Xenopus* embryos and oocytes

Adult *Xenopus laevis* wildtype J-strain females (RRID:NXR_0024) were induced to ovulate using human chorionic gonadotropin (hCG, MP Biomed.). Eggs were collected and fertilised in 0.3x Marc’s Modified Ringer’s (MMR)[1x MMR: 1 M NaCl, 18 mM KCl, 20 mM CaCl_2_,10mM MgCl_2_, and 150 mM HEPES (pH 7.6)], using a sperm suspension (J-strain wildtype or *wnt11b-/-* mutant). Embryos were dejellied at the 2-8 cell stage in 2% cysteine in 0.1x MMR (pH7.8) for 4 min before washing the embryos with 0.1x MMR. Embryos were cultured to the desired stage at 18°-24°C in 0.1x MMR. For microinjection, fertilised eggs were dejellied one hour after fertilisation as above and transferred to 2% Ficoll (Pharmacia)/0.3x MMR and injected with 2-10 nl of solution as desired using an air-driven injector (Harvard Apparatus). Injected embryos were washed into 0.1x MMR for several hours-to-overnight after injection.

Ovary was isolated from anaesthetised wild type and mutant females and divided into one centimetre segments prior to storage at 18°C. Oocytes were manually defolliculated in a modified oocyte culture medium (OCM; 67% L-15, 0.05% PVA, 1x Pen-Strep, pH 7.6-7.8; (Houston, 2018; Houston, 2019)) using watchmakers forceps (Dumont #4 or #5) and cultured at 18°C. Alternatively, for in situ hybridization, oocytes were enzymatically defolliculated by treatment with 0.02% Liberase™ (Roche Applied Science) for 1-1.5 hours in 0.5x Delbecco’s PBS with rocking, followed by extensive washing in OCM.

### CRISPR/Cas9 Mutagenesis

Mutations in *wnt11b*.*L* were generated in F0 embryos using CRISPR/Cas9 RNP injection. Guide RNAs were designed against exons 1-2 of *X. laevis* (J-strain) *wnt11b*.*L* (**Supplemental Table 1**). *Wnt11b*.*L* is the only homeolog of Wnt11b, and is the only member of the family to be expressed maternally (Session et al. 2016). In vitro transcribed guide RNAs were made from PCR-generated templates (Bhattacharya et al. 2015) and were complexed with Cas9 protein (CP01; PNA Bio) at 37°C before injection into the animal pole of fertilised eggs at the 1-2 cell stage. Mutagenesis was verified by sequencing of PCR products using DNA obtained through genotyping of whole embryo or tail samples, or through biopsy of the foot webbing of post-metamorphic animals. DNA isolation was performed as in (Bhattacharya et al., 2015) and sequencing was performed at MBL or at the Carver Center for Genomics (University of Iowa). A founder female was identified that transmitted a 13 base pair deletion in exon1 at the sgT1 site (chr8L:21220561..21220583) and was used to generate heterozygous male and female F1 offspring. These were interbred by in vitro fertilizations to generate homozygous F2 *wnt11b*.*L* mutant animals (RRID:NXR_2112). Breeding and colony maintenance were done by the NXR at MBL.

### Plasmids

Full-length cDNAs for *wnt11b*.*L* were amplified from wildtype or *wnt11b-/-* mutant oocyte total RNA using a high-fidelity polymerase (Q5, New England Biolabs). PCR products were cloned into pCR8/GW/TOPO (Invitrogen) and individual clones were verified by sequencing. Primer sequences are presented in **Supplemental Table 1**. Desired clones in the (correct) 5’L1-3’L2 orientation were inserted via recombination into a pCS2+ Gateway-converted vector (Custom vector conversion kit; Invitrogen). Details of the Gateway plasmid are available upon request. Template DNAs for sense transcripts were prepared from *wnt11b/pcs2+* plasmids by *Not*I digestion. Capped messenger RNA was synthesised using SP6 mMessage mMachine kits (Ambion). *Eb3-gfp/pcs2+* RNA was similarly prepared as described (Olson et al., 2015). Mouse *Lrp6* in pβ/RN3P was used as described (Kofron et al., 2007), prepared by *Sfi*I/T3 digestion and transcription. RNAs for zebrafish *dvl2-mcherry, rab5-gfp* and *rab11-gfp* were gifts from D. Slusarski).

### Analysis of gene expression using real-time RT-PCR

Total RNA was prepared from oocytes, embryos and explants using proteinase K and then treated with RNase-free DNase as described (Oh and Houston, 2017). Real-time RT-PCR was done using the LightCycler™ 480 system (Roche Applied Science). Samples were normalised to *ornithine decarboxylase* (*odc*) or to the geometric mean of *odc* and *fgfr1*, and relative expression values were calculated against a standard curve of control cDNA dilutions. Samples lacking reverse transcriptase in the cDNA synthesis reaction failed to give specific products. Primer sequences are listed in **Supplemental Table 1**. Charts were generated directly from the text file output of the Roche LightCycler software using a custom Python script.

### Whole-mount in situ hybridization

Whole-mount in situ hybridization was performed essentially as described (Kerr et al., 2008; Sive et al., 2000). Template DNAs for in vitro transcription were prepared by digestion, followed by transcription, with appropriate restriction enzymes and polymerases: *wnt11b/pcs2+* (*Sal*I/T7), *nodal3*.*1* (from R. Harland; *Eco*RI/T7), *eomes* and *myod1* (from J. Gurdon; *Eco*RI/T3 and *Bam*HI/SP6 respectively), *sox17a* (from A. Zorn; *Asp*718/T3), *pitx2c/pbluescript* (a gift from M. Blum; *Not*I/T7) and *sizzled/pcs2+* (from M. Kirschner, Addgene plasmid 16688, *Bam*HI/T3), *wnt8a/pcs2+* (XE10, from R. Moon; Addgene plasmid 16865; *Bam*HI/T3). Antisense RNA probes labelled with digoxygenin-11-UTP (Roche) were synthesised using polymerases and reaction buffers from Promega. Processing of in situ hybridization was performed manually or using a robotic system (Biolane, Intavis AG).

### Immunoblotting

Samples were lysed in cell lysis buffer (150 mM NaCl, 20 mM Tris-HCl (pH 7.4), 1 mM EDTA, 1 mM EGTA (pH 8.0), 1% Triton X-100, with protease and/or phosphatase inhibitors) and clarified by centrifugation (10 minutes at 10,000 x g). Lysates were heated in sample buffer (LAEMMLI, 1970) and the equivalent of 0.5 - 3 oocytes/embryos were electrophoresed on SDS-PAGE TGX AnyKD Ready Gels (BioRad) or large homemade gels and transferred to nitrocellulose (Power Blotter, Thermo-Pierce). For anti-phospho-LRP6 analysis, samples were lysed fresh (without freezing), and anti-phospho-LRP6 was used first, before stripping and reblotting.

Membranes were blocked in 5% BSA in TBS, 0.1% Tween 20 (for anti-phospho-LRP6) or 5% nonfat dry milk (HyVee) in PBS, 0.1% Tween 20, and incubated in primary antibody overnight at 4°C. Detection was performed using mouse or rabbit secondary antibodies conjugated to peroxidase (1:10000, Jackson Immunoresearch), and Licor reagents and equipment (C-Digit Scanner). Antibodies and dilutions used were: rabbit anti-phospho-LRP6 polyclonal antibodies (S1490; 1:500; Cell Signaling Technology (CST); RRID:AB_2139327), anti-LRP6 rabbit mAbs (C5C7 and C47E12; 1:500 each, mixed together; CST; RRID:AB_2139329 and RRID:AB_1950408), anti-Beta-tubulin (mAb E7; 1:1000; DSHB; RRID:AB_528499), and diphospho-ERK-1 and ERK-2 (1:4000; clone MAPK-YT; Sigma; RRID:AB_477245). In our hands, anti-phospho-LRP6 recognises endogenous *Xenopus* and injected mouse phospho-S1490 Lrp6, whereas the non-phospho mAbs recognise mouse but not frog Lrp6.

### Immunostaining

Whole-mount immunostaining was performed on embryos fixed in MEMFA and stored in 100% methanol (embryo stages) or fixed directly in methanol (eggs), using modifications of previously described methods (Cuykendall and Houston, 2009; Elinson and Rowning, 1988). Fixed samples were rehydrated gradually to 1x PBS, then to PBT(PBS/0.5% Triton X-100/0.2% BSA (fraction V)) and then blocked for two hours at room temperature in PBT/2.5% BSA. Samples were washed for one hour the next day in PBT and incubated with primary antibodies diluted in PBT overnight at 4°C with rocking, followed by five one hour washes in PBT. Incubation with secondary antibodies diluted in PBT was done for 2 hrs followed by washing as above.

Antibodies were anti-Beta-tubulin mAb E7 (1:200 dilution of monoclonal antibody supernatant concentrate; DSHB Hybridoma Product E7, deposited with the DSHB by M. Klymkowsky; (Chu and Klymkowsky, 1989); RRID:AB_528499). Secondary antibodies were goat anti-mouse Alexa-488, diluted in PBT (1:500; Invitrogen/Molecular Probes). Fluorescence was visualised on a Leica DMI4000B inverted microscope using 20X - 63X dry objectives (Leica Microsystems). For confocal analysis, samples were imaged on an SP8 confocal imaging system (Leica Microsystems) using a 20X objective with or without tile-scanning to visualise the entire vegetal surface.

### Time-lapse image analysis of microtubule plus end dynamics

Samples were imaged at room temperature on an inverted, wide-field epi fluorescence microscope (DMI4000B, Leica Microsystems) using an oil-immersion Leica 100x /1.30 N.A. PLANAPO objective. Image acquisition was done using a Leica DFC3000G monochrome camera at a frame rate of two seconds per frame and using Leica LAS software. The pixel size was 0.0536 × 0.0536 × 0.200 mm and the image size was 1296 × 966 pixels. Leica files (.lif) were imported directly into MATLAB_2021a using u-track (v2.3; https://github.com/DanuserLab/u-track; (Applegate et al., 2011; Jaqaman et al., 2008; Matov et al., 2010)) and processed using similar parameters as described (Olson et al., 2015). Custom code was inserted to incorporate code from the CircStat circular statistics toolbox (Berens, 2009) into the u-track analysis (available upon request). Statistical analyses of these data were performed using MATLAB; the fdr_bh package was used for adjusted p-values (Groppe, 2022; (Benjamini, 2010; Benjamini and Hochberg, 1995)). Consecutive frame averaging was performed using the Time-lapse Series Painter for FIJI (Vaart et al., 2011). Images for publication were prepared in Adobe Illustrator and Photoshop using only level and contrast adjustments applied over the entire image. Other modifications included resizing, changing stroke/fill weights and colours and annotated overlays.

## Acknowledgements

The authors would like to thank Cindy Toll for technical assistance in library preparation. We also thank Profs. R. Harland, J. Gurdon, A. Zorn, M. Kirschner, M. Blum, R. Lang, D. Slusarski, and R. Moon for contributing reagents (either directly or through Addgene and DSHB). We acknowledge Xenbase.org, the National Xenopus Resource, Addgene, and the Developmental Studies Hybridoma Bank (DSHB) for critical community resources. This work was funded by the University of Iowa (D.W.H) and by grants from the NIH, R01GM083999 (D.W.H) and R24OD030008, P40OD010997 (M.E.H.).

## Data availability

GEO accession GSE195806.

## References

Applegate, K. T., Besson, S., Matov, A., Bagonis, M. H., Jaqaman, K. and Danuser, G. (2011). plusTipTracker: Quantitative image analysis software for the measurement of microtubule dynamics. Journal of structural biology 176, 168–184.

Benjamini, Y. (2010). Discovering the false discovery rate. J Royal Statistical Soc Ser B Statistical Methodol 72, 405–416.

Benjamini, Y. and Hochberg, Y. (1995). Controlling the False Discovery Rate: A Practical and Powerful Approach to Multiple Testing. J Royal Statistical Soc Ser B Methodol 57, 289–300.

Berens, P. (2009). CircStat: a MATLAB toolbox for circular statistics. J Stat Softw 31, 1–21.

Bhattacharya, D., Marfo, C. A., Li, D., Lane, M. and Khokha, M. K. (2015). CRISPR/Cas9: An inexpensive, efficient loss of function tool to screen human disease genes in Xenopus. Developmental Biology 408, 196–204.

Bilic, J., Huang, Y.-L., Davidson, G., Zimmermann, T., Cruciat, C.-M., Bienz, M. and Niehrs, C. (2007). Wnt induces LRP6 signalosomes and promotes dishevelled-dependent LRP6 phosphorylation. Science 316, 1619–1622.

Blythe, S. A., Cha, S.-W., Tadjuidje, E., Heasman, J. and Klein, P. S. (2010). beta-Catenin primes organizer gene expression by recruiting a histone H3 arginine 8 methyltransferase, Prmt2. Developmental cell 19, 220–231.

Bolce, M. E., Hemmati-Brivanlou, A., Kushner, P. D. and Harland, R. M. (1992). Ventral ectoderm of Xenopus forms neural tissue, including hindbrain, in response to activin. Dev Camb Engl 115, 681–8.

Cha, S.-W., Tadjuidje, E., Tao, Q., Wylie, C. C. and Heasman, J. (2008). Wnt5a and Wnt11 interact in a maternal Dkk1-regulated fashion to activate both canonical and non-canonical signaling in Xenopus axis formation. Development (Cambridge, England) 135, 3719–3729.

Chan, A. P., Kloc, M., Larabell, C. A., LeGros, M. and Etkin, L. D. (2007). The maternally localized RNA fatvg is required for cortical rotation and germ cell formation. Mechanisms of development 124, 350–363.

Chu, D. T. and Klymkowsky, M. W. (1989). The appearance of acetylated alpha-tubulin during early development and cellular differentiation in Xenopus. Developmental Biology 136, 104–117.

Cuykendall, T. N. and Houston, D. W. (2009). Vegetally localized Xenopus trim36 regulates cortical rotation and dorsal axis formation. Development 136, 3057–3065.

Darras, S., Marikawa, Y., Elinson, R. P. and Lemaire, P. (1997). Animal and vegetal pole cells of early Xenopus embryos respond differently to maternal dorsal determinants: implications for the patterning of the organiser. Development (Cambridge, England) 124, 4275–4286.

Dichmann, D. S., Walentek, P. and Harland, R. M. (2015). The Alternative Splicing Regulator Tra2b Is Required for Somitogenesis and Regulates Splicing of an Inhibitory Wnt11b Isoform. Cell Reports 10, 527–536.

Dobrowolski, R. and Robertis, E. M. D. (2012). Endocytic control of growth factor signalling: multivesicular bodies as signalling organelles. Nat Rev Mol Cell Bio 13, 53–60.

Domenico, E. D., Owens, N. D. L., Grant, I. M., Gomes-Faria, R. and Gilchrist, M. J. (2015). Molecular asymmetry in the 8-cell stage Xenopus tropicalis embryo described by single blastomere transcript sequencing. Dev Biol 408, 252–268.

Du, S. J., Purcell, S. M., Christian, J. L., McGrew, L. L. and Moon, R. T. (1995). Identification of distinct classes and functional domains of Wnts through expression of wild-type and chimeric proteins in Xenopus embryos. Molecular and cellular biology 15, 2625–2634.

Dyle, M. C., Kolakada, D., Cortazar, M. A. and Jagannathan, S. (2020). How to get away with nonsense: Mechanisms and consequences of escape from nonsense-mediated RNA decay. Wiley Interdiscip Rev Rna 11, e1560.

Dzamba, B. J., Jakab, K. R., Marsden, M., Schwartz, M. A. and DeSimone, D. W. (2009). Cadherin Adhesion, Tissue Tension, and Noncanonical Wnt Signaling Regulate Fibronectin Matrix Organization. Dev Cell 16, 421–432.

El-Brolosy, M. A., Kontarakis, Z., Rossi, A., Kuenne, C., Günther, S., Fukuda, N., Kikhi, K., Boezio, G. L. M., Takacs, C. M., Lai, S.-L., et al. (2019). Genetic compensation triggered by mutant mRNA degradation. Nature 568, 193–197.

Elinson, R. P. and Rowning, B. (1988). A transient array of parallel microtubules in frog eggs: potential tracks for a cytoplasmic rotation that specifies the dorso-ventral axis. Developmental Biology 128, 185–197.

Flachsova, M., Sindelka, R. and Kubista, M. (2013). Single blastomere expression profiling of Xenopus laevis embryos of 8 to 32-cells reveals developmental asymmetry. Scientific Reports 3,.

Garriock, R. J., D’Agostino, S. L., Pilcher, K. C. and Krieg, P. A. (2005). Wnt11-R, a protein closely related to mammalian Wnt11, is required for heart morphogenesis in Xenopus. Developmental Biology 279, 179–192.

Gerhart, J. (2004). Symmetry breaking in the egg of Xenopus laevis.pp. 341–351.

Heisenberg, C.-P., Tada, M., Rauch, G.-J., Saúde, L., Concha, M. L., Geisler, R., Stemple, D. L., Smith, J. C. and Wilson, S. W. (2000). Silberblick/Wnt11 mediates convergent extension movements during zebrafish gastrulation. Nature 405, 76–81.

Hino, H., Nakanishi, A., Seki, R., Aoki, T., Yamaha, E., Kawahara, A., Shimizu, T. and Hibi, M. (2018). Roles of maternal wnt8a transcripts in axis formation in zebrafish. Developmental Biology 434, 96–107.

Hoppler, S., Brown, J. D. and Moon, R. T. (1996). Expression of a dominant-negative Wnt blocks induction of MyoD in Xenopus embryos. Gene Dev 10, 2805–2817.

Houston, D. W. (2012). Cortical rotation and messenger RNA localization in Xenopus axis formation. Wiley Interdiscip Rev Dev Biology 1, 371–388.

Houston, D. W. (2013). Regulation of cell polarity and RNA localization in vertebrate oocytes. International review of cell and molecular biology 306, 127–185.

Houston, D. W. (2017). Vertebrate axial patterning: from egg to asymmetry. In Vertebrate Development, Maternal to Zygotic Control (ed. Pelegri, F.), Sutherland, A.), and Danilchik, M.), pp. 209–306. Cham: Springer.

Houston, D. W. (2018). Oocyte Host-Transfer and Maternal mRNA Depletion Experiments in Xenopus. Cold Spring Harbor protocols 10, pdb–prot096982.

Houston, D. W. (2019). Culture and Host Transfer of Xenopus Oocytes for Maternal mRNA Depletion and Genome Editing Experiments. In (ed. Pelegri, F.), pp. 1–16. New York: Humana.

Huebner, R. J. and Wallingford, J. B. (2018). Coming to Consensus: A Unifying Model Emerges for Convergent Extension. Dev Cell 46, 389–396.

Jaqaman, K., Loerke, D., Mettlen, M., Kuwata, H., Grinstein, S., Schmid, S. L. and Danuser, G. (2008). Robust single particle tracking in live cell time-lapse sequences. Nat Methods 5, 695–702.

Jussila, M. and Ciruna, B. (2017). Zebrafish models of non-canonical Wnt/planar cell polarity signalling: fishing for valuable insight into vertebrate polarized cell behavior. Wiley Interdiscip Rev Dev Biology 6,.

Kageura, H. (1997). Activation of dorsal development by contact between the cortical dorsal determinant and the equatorial core cytoplasm in eggs of Xenopus laevis. Dev Camb Engl 124, 1543–51.

Kao, K. R. and Elinson, R. P. (1988). The entire mesodermal mantle behaves as Spemann’s organizer in dorsoanterior enhanced Xenopus laevis embryos. Developmental Biology 127, 64–77.

Kerr, T. C., Cuykendall, T. N., Luettjohann, L. C. and Houston, D. W. (2008). Maternal Tgif1 regulates nodal gene expression in Xenopus. Developmental dynamics 237, 2862–2873.

Kim, S.-E., Huang, H., Zhao, M., Zhang, X., Zhang, A., Semonov, M. V., MacDonald, B. T., Zhang, X., Abreu, J. G., Peng, L., et al. (2013). Wnt Stabilization of β-Catenin Reveals Principles for Morphogen Receptor-Scaffold Assemblies. Science 340, 867–870.

Kintner, C. R. and Brockes, J. P. (1984). Monoclonal antibodies identify blastemal cells derived from dedifferentiating limb regeneration. Nature 308, 67–9.

Kofron, M., Birsoy, B., Houston, D., Tao, Q., Wylie, C. and Heasman, J. (2007). Wnt11/β-catenin signaling in both oocytes and early embryos acts through LRP6-mediated regulation of axin. Development 134, 503–513.

Kraft, B., Berger, C. D., Wallkamm, V., Steinbeisser, H. and Wedlich, D. (2012). Wnt-11 and Fz7 reduce cell adhesion in convergent extension by sequestration of PAPC and C-cadherin. The Journal of cell biology 198, 695–709.

Kushner, P. D. (1984). A Library of Monoclonal Antibodies to Torpedo Cholinergic Synaptosomes. J Neurochem 43, 775–786.

Laemmli, U. K. (1970). Cleavage of Structural Proteins during the Assembly of the Head of Bacteriophage T4. Nature 227, 680–685.

Lamb, T. M., Knecht, A. K., Smith, W. C., Stachel, S. E., Economides, A. N., Stahl, N., Yancopolous, G. D. and Harland, R. M. (1993). Neural Induction by the Secreted Polypeptide Noggin. Science 262, 713–718.

Leyns, L., Bouwmeester, T., Kim, S. H., Piccolo, S. and Robertis, E. M. D. (1997). Frzb-1 is a secreted antagonist of Wnt signaling expressed in the Spemann organizer. Cell 88, 747–756.

Li, V. S. W., Ng, S. S., Boersema, P. J., Low, T. Y., Karthaus, W. R., Gerlach, J. P., Mohammed, S., Heck, A. J. R., Maurice, M. M., Mahmoudi, T., et al. (2012). Wnt Signaling through Inhibition of β-Catenin Degradation in an Intact Axin1 Complex. Cell 149, 1245–1256.

Ma, Z., Zhu, P., Shi, H., Guo, L., Zhang, Q., Chen, Y., Chen, S., Zhang, Z., Peng, J. and Chen, J. (2019). PTC-bearing mRNA elicits a genetic compensation response via Upf3a and COMPASS components. Nature 568, 259–263.

Ma, W., Chen, M., Kang, H., Steinhart, Z., Angers, S., He, X. and Kirschner, M. W. (2020). Single-molecule dynamics of Dishevelled at the plasma membrane and Wnt pathway activation. P Natl Acad Sci Usa 117, 16690–16701.

Marikawa, Y. and Elinson, R. P. (1999). Relationship of vegetal cortical dorsal factors in the Xenopus egg with the Wnt/beta-catenin signaling pathway. Mechanisms of development 89, 93–102.

Marikawa, Y., Li, Y. and Elinson, R. P. (1997). Dorsal determinants in the Xenopus egg are firmly associated with the vegetal cortex and behave like activators of the Wnt pathway. Developmental Biology 191, 69–79.

Matov, A., Applegate, K., Kumar, P., Thoma, C., Krek, W., Danuser, G. and Wittmann, T. (2010). Analysis of microtubule dynamic instability using a plus-end growth marker. Nature methods 7, 761–768.

Miller, J. R., Rowning, B. A., Larabell, C. A., Yang-Snyder, J. A., Bates, R. L. and Moon, R. T. (1999). Establishment of the Dorsal–Ventral Axis inXenopus Embryos Coincides with the Dorsal Enrichment of Dishevelled That Is Dependent on Cortical Rotation. The Journal of Cell Biology 146, 427–438.

Myers, D. C., Sepich, D. S. and Solnica-Krezel, L. (2002). Bmp activity gradient regulates convergent extension during zebrafish gastrulation. Developmental Biology 243, 81–98.

Nonaka, S., Tanaka, Y., Okada, Y., Takeda, S., Harada, A., Kanai, Y., Kido, M. and Hirokawa, N. (1998). Randomization of Left–Right Asymmetry due to Loss of Nodal Cilia Generating Leftward Flow of Extraembryonic Fluid in Mice Lacking KIF3B Motor Protein. Cell 95, 829–837.

Oh, D. and Houston, D. W. (2017). Role of maternal Xenopus syntabulin in germ plasm aggregation and primordial germ cell specification. Developmental Biology 432, 237–247.

Pearl, E. J., Grainger, R. M., Guille, M. and Horb, M. E. (2012). Development of xenopus resource centers: The national xenopus resource and the european xenopus resource center. Genesis 50, 155–163.

Rim, E. Y., Kinney, L. K. and Nusse, R. (2020). β-catenin-mediated Wnt signal transduction proceeds through an endocytosis-independent mechanism. Mol Biol Cell 31, 1425–1436.

Schweickert, A., Weber, T., Beyer, T., Vick, P., Bogusch, S., Feistel, K. and Blum, M. (2007). Cilia-driven leftward flow determines laterality in Xenopus. Current Biology 17, 60–66.

Schweickert, A., Vick, P., Getwan, M., Weber, T., Schneider, I., Eberhardt, M., Beyer, T., Pachur, A. and Blum, M. (2010). The Nodal Inhibitor Coco Is a Critical Target of Leftward Flow in Xenopus. Current Biology.

Session, A. M., Uno, Y., Kwon, T., Chapman, J. A., Toyoda, A., Takahashi, S., Fukui, A., Hikosaka, A., Suzuki, A., Kondo, M., et al. (2016). Genome evolution in the allotetraploid frog Xenopus laevis. Nature 538, 336–343.

Sive, H. L., Grainger, R. M. and Harland, R. M. (2000). Early development of Xenopus laevis: a laboratory manual.

Solnica-Krezel, L. (2005). Conserved patterns of cell movements during vertebrate gastrulation. Current Biology 15, R213–28.

Tamai, K., Zeng, X., Liu, C., Zhang, X., Harada, Y., Chang, Z. and He, X. (2004). A mechanism for Wnt coreceptor activation. Molecular cell 13, 149–156.

Tao, Q., Yokota, C., Puck, H., Kofron, M., Birsoy, B., Yan, D., Asashima, M., Wylie, C. C., Lin, X. and Heasman, J. (2005). Maternal Wnt11 Activates the Canonical Wnt Signaling Pathway Required for Axis Formation in Xenopus Embryos. Cell 120, 857–871.

Tochinai, S. and Katagiri, C. (1975). COMPLETE ABROGATION OF IMMUNE RESPONSE TO SKIN ALLOGRAFTS AND RABBIT ERYTHROCYTES IN THE EARLY THYMECTOMIZED XENOPUS. Dev Growth Differ 17, 383–394.

Ulrich, F., Concha, M., Heid, P., Voss, E., Witzel, S., Roehl, H., Tada, M., Wilson, S., Adams, R. J., Soll, D., et al. (2003). Slb/Wnt11 controls hypoblast cell migration and morphogenesis at the onset of zebrafish gastrulation. Development (Cambridge, England) 130, 5375–5384.

Ulrich, F., Krieg, M., Schötz, E.-M., Link, V., Castanon, I., Schnabel, V., Taubenberger, A., Mueller, D., Puech, P.-H. and Heisenberg, C.-P. (2005). Wnt11 Functions in Gastrulation by Controlling Cell Cohesion through Rab5c and E-Cadherin. Dev Cell 9, 555–564.

Vaart, B. van der, Manatschal, C., Grigoriev, I., Olieric, V., Gouveia, S. M., Bjelić, S., Demmers, J., Vorobjev, I., Hoogenraad, C. C., Steinmetz, M. O., et al. (2011). SLAIN2 links microtubule plus end–tracking proteins and controls microtubule growth in interphase. J Cell Biol 193, 1083–1099.

Van Itallie, E. S., Field, C. M., Mitchison, T. J. and Kirschner, M. W. (2022). Wnt11 family dependent morphogenesis during frog gastrulation is marked by the cleavage furrow protein anillin. Biorxiv 2022.01.07.475368.

Walentek, P., Schneider, I., Schweickert, A. and Blum, M. (2013). Wnt11b Is Involved in Cilia-Mediated Symmetry Breakage during Xenopus Left-Right Development. Plos One 8, e73646.

Wang, S., Krinks, M., Lin, K., Luyten, F. and Moos, M. (1997). Frzb, a secreted protein expressed in the Spemann organizer, binds and inhibits Wnt-8. Cell 88, 757–766.

Weaver, C. and Kimelman, D. (2004). Move it or lose it: axis specification in Xenopus. Development (Cambridge, England) 131, 3491–3499.

Weaver, C., Farr, G. H., Pan, W., Rowning, B. A., Wang, J., Mao, J., Wu, D., Li, L., Larabell, C. A. and Kimelman, D. (2003). GBP binds kinesin light chain and translocates during cortical rotation in Xenopus eggs. Development (Cambridge, England) 130, 5425–5436.

Wen, J. W. and Winklbauer, R. (2017). Ingression-type cell migration drives vegetal endoderm internalisation in the Xenopus gastrula. Elife 6, e27190.

Winklbauer, R. and Schürfeld, M. (1999). Vegetal rotation, a new gastrulation movement involved in the internalization of the mesoderm and endoderm in Xenopus. Development (Cambridge, England) 126, 3703–3713.

Winklbauer, R., Medina, A., Swain, R. K. and Steinbeisser, H. (2001). Frizzled-7 signalling controls tissue separation during Xenopus gastrulation. Nature 413, 856–860.

Witzel, S., Zimyanin, V., Carreira-Barbosa, F., Tada, M. and Heisenberg, C.-P. (2006). Wnt11 controls cell contact persistence by local accumulation of Frizzled 7 at the plasma membrane. J Cell Biology 175, 791–802.

Xing, Y.-Y., Cheng, X.-N., Li, Y.-L., Zhang, C., Saquet, A., Liu, Y.-Y., Shao, M. and Shi, D.-L. (2018). Mutational analysis of dishevelled genes in zebrafish reveals distinct functions in embryonic patterning and gastrulation cell movements. Plos Genet 14, e1007551.

Yan, L., Chen, J., Zhu, X., Sun, J., Wu, X., Shen, W., Zhang, W., Tao, Q. and Meng, A. (2018). Maternal Huluwa dictates the embryonic body axis through β-catenin in vertebrates. Science 362, eaat1045.

Yang, J., Tan, C., Darken, R. S., Wilson, P. A. and Klein, P. S. (2002). Beta-catenin/Tcf-regulated transcription prior to the midblastula transition. Development (Cambridge, England) 129, 5743–5752.

Yokota, C., Kofron, M., Zuck, M., Houston, D. W., Isaacs, H., Asashima, M., Wylie, C. C. and Heasman, J. 2003). A novel role for a nodal-related protein; Xnr3 regulates convergent extension movements via the FGF receptor. Development (Cambridge, England) 130, 2199–2212.

Yost, H. J. (1992). Regulation of vertebrate left–right asymmetries by extracellular matrix. Nature 357, 158–161.

Zeng, X., Tamai, K., Doble, B., Li, S., Huang, H., Habas, R., Okamura, H., Woodgett, J. and He, X. (2005). A dual-kinase mechanism for Wnt co-receptor phosphorylation and activation. Nature 438, 873–877.

